# Nanopore Long-Read Sequencing Unveils Genomic Disruptions in Alzheimer’s Disease

**DOI:** 10.1101/2024.02.01.578450

**Authors:** Paulino Ramirez, Wenyan Sun, Shiva Kazempour Dehkordi, Habil Zare, Giovanni Pascarella, Piero Carninci, Bernard Fongang, Kevin F. Bieniek, Bess Frost

## Abstract

Studies in laboratory models and postmortem human brain tissue from patients with Alzheimer’s disease have revealed disruption of basic cellular processes such as DNA repair and epigenetic control as drivers of neurodegeneration. While genomic alterations in regions of the genome that are rich in repetitive sequences, often termed “dark regions,” are difficult to resolve using traditional sequencing approaches, long-read technologies offer promising new avenues to explore previously inaccessible regions of the genome. In the current study, we leverage nanopore-based long-read whole-genome sequencing of DNA extracted from postmortem human frontal cortex at early and late stages of Alzheimer’s disease, as well as age-matched controls, to analyze retrotransposon insertion events, non-allelic homologous recombination (NAHR), structural variants and DNA methylation within retrotransposon loci and other repetitive/dark regions of the human genome. Interestingly, we find that retrotransposon insertion events and repetitive element-associated NAHR are particularly enriched within centromeric and pericentromeric regions of DNA in the aged human brain, and that ribosomal DNA (rDNA) is subject to a high degree of NAHR compared to other regions of the genome. We detect a trending increase in potential somatic retrotransposition events of the small interfering nuclear element (SINE) AluY in late-stage Alzheimer’s disease, and differential changes in methylation within repetitive elements and retrotransposons according to disease stage. Taken together, our analysis provides the first long-read DNA sequencing-based analysis of retrotransposon sequences, NAHR, structural variants, and DNA methylation in the aged brain, and points toward transposable elements, centromeric/pericentromeric regions and rDNA as hotspots for genomic variation.

## INTRODUCTION

The brain accumulates structural variants that result in genomic mosaicism between cells over the course of human aging [1]. Structural variants can occur in progenitor cells during development, proliferating glial and epithelial cells, and in post-mitotic neurons. Such genetic variants can arise from biological processes such as DNA repair, homologous recombination, replication and retrotransposition. Studies of the human brain report changes in DNA content, DNA copy number variation, somatic recombination, and tandem repeat expansions, revealing a prevalent accumulation of genomic rearrangements [2-4]. These genomic changes have also been reported in neurodegenerative diseases, including Alzheimer’s disease, for which aging is the largest risk factor [5]. The functional impacts of various structural variants depend on the cell type in which they occur and the region of the genome affected.

Class II transposable elements, the “retrotransposons,” compose 35% of the human genome [6] and are a potential source of genomic variation. Active retrotransposons follow a life cycle in which mRNA is reverse transcribed into complementary DNA that can then be integrated back into the host genome. Some retrotransposons encode protein products such as reverse transcriptase and endonuclease that facilitate retrotransposition [7]. In the human genome, specific subfamilies of long interspersed element (LINE) retrotransposons are capable of autonomous retrotransposition, while some active SINE and SINE-VNTR-Alu (SVA) retrotransposons leverage LINE-encoded protein for trans-retrotransposition. Retrotransposition of the active human-specific LINE-1 element (L1Hs) has been documented in neurons and glia and is proposed to be a natural occurrence during neurodevelopment [8-10]. Somatic retrotransposition has been reported in the context of neurodegeneration in various laboratory models of disease [11], but has not been studied in the aged human brain. In addition to *de novo* retrotransposition events, the repetitive nature of retrotransposon DNA loci has the potential to facilitate NAHR, leading to the generation of new genomic variants [12-16]. These events, which have been documented in neurodegenerative diseases, arise from double stranded DNA breaks that occur during instances of high cellular stress and can contribute to the loss of genomic integrity [17].

The advancement of long-read sequencing now allows the resolution of “dark regions” (a term coined by Ebbert et al., 2019), highly repetitive regions of the genome that are difficult to sequence using traditional approaches [18-21]. These genomic stretches include low-complexity regions consisting of simple and tandem repeats, as well as repetitive regions rich in transposable elements. Traditionally, dark regions have been difficult to characterize due to either lack of sequencing depth or short-read associated mapping quality, with many of these loci originating from genomic duplication [18]. Long-read sequencing can generate reads spanning an entire structural variant and flanking regions, allowing for more confident identification of variants present in dark regions [18, 22, 23]. This extends to the proper identification and quantification of germline-derived and somatic retrotransposon insertions, including those present in only a single cell [24], and NAHR between repetitive elements. While many dark regions are present in genes previously implicated in human disease, our inability to resolve their sequences with affordable high-throughput technologies has led to their exclusion from analysis in most studies.

The combination of long reads and accompanying calling of 5-methylcytosine (5mC) methylation sites allow for more comprehensive analysis of low complexity and repetitive regions [25] that are generally ignored in classical methylation analysis methods [26]. Overall, genomic CpG methylation tends to decline as organisms grow older [27], particularly in various human age-related conditions such as cancer [28] and Alzheimer’s disease [29-34]. The use of long-read methylation data can further complement previous analyses by accounting for methylation of previously discarded dark regions.

In the current study, we utilize Oxford Nanopore DNA sequencing to analyze putatively somatic retrotransposon insertions, NAHR between repetitive elements, structural variants, and DNA methylation in frontal cortex samples from aged individuals affected by Alzheimer’s disease. Neuropathological changes in the aged human brain are common; most individuals over the age of 60 have some degree of deposition of pathological forms of tau [35]. Tau protein accumulates in a diverse group of neurodegenerative disorders, including Alzheimer’s disease, that are collectively referred to as “tauopathies.” Alzheimer’s disease is the most common neurodegenerative disorder, with an incidence of 5% of individuals aged 65-74, 13.1% of individuals aged 75-84, and 33.2% of individuals aged 85+ [36]. The pathological distribution of tau in Alzheimer’s disease is defined by Braak neurofibrillary tangle staging. Compared to a brain void of tau pathology (Stage 0), tau pathology progresses from brainstem structures (Stages a-c) to transentorhinal/entorhinal regions (Stage I), to the primary hippocampus (Stage III) and ultimately, association and primary neocortical regions of the brain (Stages V-VI) [37]. To capture changes related to the neuropathological progression of tau pathology in Alzheimer’s disease, we included equal numbers of control (Braak 0), mid-stage (Braak III), and late-stage (Braak V/VI) cases among the 18 frontal cortex samples analyzed.

Our analyses reveal that repetitive segments of the genome are particularly susceptible to genomic changes in the aged human brain. Centromeric and pericentromeric regions of chromosomes are enriched with retrotransposon insertions and NAHR events between repetitive elements. Additionally, 47S rDNA exhibits a particularly high frequency of NAHR compared to other genomic regions. We also observe a trend toward increased putatively somatic retrotransposition events involving the SINE AluY family in advanced stages of Alzheimer’s disease, alongside stage-specific differences in methylation patterns within repetitive elements and retrotransposons. Overall, this study represents the first long-read DNA sequencing-based investigation of retrotransposons, NAHR, structural variants, and DNA methylation in the aged brain, identifying transposable elements, centromeric/pericentromeric regions, and rDNA as key hotspots of genomic variation.

## RESULTS

### Nanopore long-read sequencing of DNA extracted from aged human brain

The 18 human frontal cortex samples utilized in this study had an average age of 76.6 years (**Table 1**). DNA was extracted from isolated nuclei of postmortem frontal cortex of six individuals with no clinical or pathological diagnosis of neurodegeneration (Braak 0), six individuals at Braak stage III and six individuals at Braak stage V/VI with a clinical diagnosis of Alzheimer’s disease (**Supplemental Table 1**). Sequencing was performed using the Oxford Nanopore Promethion sequencing platform. The mean read quality score was 11, with a n50 of 23.2 kbp and an average genomic coverage of 7.38X (**Supplemental Table 2**).

**Table 1.** Brain demographics and sequencing metrics. Sample characteristics and associated sequencing statistics.

|  | Braak 0 | Braak III | Braak V/VI | All | Range |
| --- | --- | --- | --- | --- | --- |
| Total # sequenced | 6 | 6 | 6 | 18 |  |
| Age | 76 | 82.3 | 71.3 | 76.6 | 67-92 |
| % Female | 50% | 50% | 33% | 44% |  |
| Postmortem interval | 10.33 h | 14.67 h | 8.75 h | 11.89 h | 5-24 |

|  | Mean | Range |
| --- | --- | --- |
| Total bases sequenced | 25.5 gbp | 14.2-41.6 gbp |
| Read quality | 11.7 | 11.5-11.9 |
| Read length (kbp) | 23.2 kbp | 9.66-31.58 kbp |
| Genome coverage | 7.4X | 4.1X-12.1X |

### Detection of retrotransposon singleton insertions in the aged human brain

We first analyzed non-reference genome retrotransposons across all 18 human brain samples using TLDR [38], a tool designed for long-read sequencing data to detect retrotransposon insertions that are lacking in the current human reference genome assemblies (the new telomere-to-telomere CHM13 and previous GRCh38 genome). These non-reference retrotransposons reflect retrotransposons that are polymorphic within the human genome that the individuals used to generate reference genomes happened to lack, and retrotransposition events that occurred in a given individual *in utero* or during life. Among the 18 human brains analyzed, we detect 642 non-reference retrotransposon sequences that are unique to a single sample and are not present in the CHM13 or GRCh38 genomes (**Supplemental Fig. 1A**).

We next identified potential somatic retrotransposition events marked by singleton insertions (insertions only supported by a single read) following methods similar to Siudeja and colleagues [24]. As the site of L1-mediated retrotransposon insertion is random, *de novo* insertions should be present as singletons if they occur in post-mitotic cells, or are present at low frequency compared to germline insertions if they occur in dividing cells. Of all non-reference insertions unique to a given individual, 54 were identified as singletons, had at least ten additional reads covering the location of the insertion, all of which lacked the insertion, and possessed target site duplication (TSD)-like flanking sequences indicative of L1-mediated transposition (**Supplemental File 1**). Intact TSDs are a signature of a recent target-primed reverse transcription event that consists of a duplication of the target site motif (canonically AA/TTTT) of insertion on either side of the inserted element [39-43]. Among retrotransposon families, the Alu subfamily of SINE elements is the most abundant class of TSD-harboring singleton insertions in the aged human brain; we find that AluYa5 is the most active member within the Alu subfamily (**Fig. 1A**), consistent with previous work in humans [44]. Interestingly, we find that the majority of singleton insertions (unlike non-singleton insertions) are located in centromeric and pericentromeric regions of the genome, predominantly occurring in chromosomes 11, 14, 16, and 20 (**Supplemental Fig. 1B**). Our findings align with previous research indicating that centromeres may function as sinks for retrotransposon insertions [45-47]. Further analysis of singleton Alu insertions with respect to centromeric repeats (**Fig. 1B**) reveals that they are predominantly located in alpha satellite active highly ordered repeat (HOR) regions, while singleton L1 insertions are more frequent in inactive HORs **(Fig. 1C)**. Functionally, active HORs serve as binding sites for kinetochore proteins during cell division.

**Figure 1.**
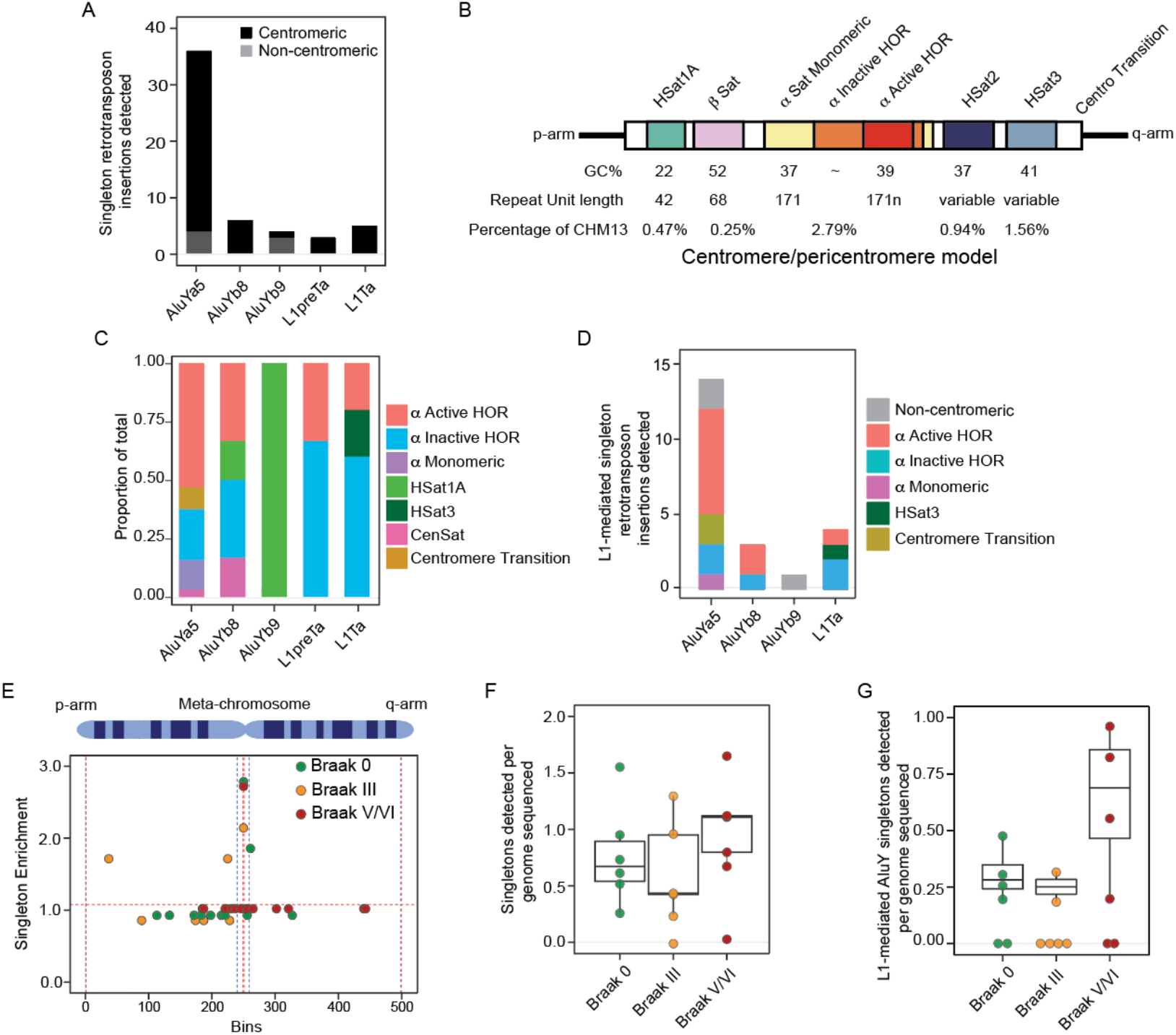
Enrichment of singleton retrotransposon insertions in the centromere of the aged human brain. A) Total counts of all singleton retrotransposon insertions by subfamily grouped by whether they are present in the centromere. B) Graphic model of centromeric and pericentromic regions. C) Barplot of the fraction of singleton insertions detected within functional centromeric repeats. D) Total counts of all L1en-mediated singleton retrotransposon insertions by subfamily, grouped by centromeric repeat. E) Mean count of retrotransposon singleton insertions detected across all autosomes, normalized for chromosome length plotted across a meta-chromosome. F) Boxplots of singleton retrotransposon insertion counts normalized to sequencing depth. G) Boxplots of L1 endonuclease-like mediated singleton AluY insertion counts normalized to sequencing depth.

Some LINE and SINE retrotransposons are transpositionally active in somatic cells, including postmitotic neurons [48-51]. Using previously established criteria, we next characterized L1 endonuclease-mediated (L1en) insertion events from evolutionarily “young” elements by filtering our list of singleton insertion events to include reads of 10 kbp or longer that harbor characteristics of L1en transposition, including presence of a polyA tail and an L1 endonuclease motif at the predicted insertion location. 22 putative L1en singleton insertions were detected across the 18 brains analyzed, most of which belonged to the AluYa5 subfamily (**Fig. 1D**). While most Alu insertions are intact, L1en-mediated LINE-1 singleton insertions are severely truncated at the 5’ ends of the L1Hs element, consistent with a 3’ end bias of insertion [52] (**Supplemental Table 1**). We attempted to determine the origin of L1Hs singleton insertions via Blat alignment of the insertion sequences against the CHM13 genome, but alignments were ambiguous due to lack of transduction sequences. In line with unique singletons, we find significant enrichment (P value of 1E-15, Fisher-Test) of L1en insertions in centromeric regions of the genome, with 17 occurring in HOR segments of the centromere. Enrichment of L1en singleton insertions in the centromere compared to all unique insertions and general singletons suggests that somatic retrotransposition events occur more commonly in the centromere.

Retrotransposon transcripts, particularly those of the endogenous retrovirus (ERV) class, are significantly elevated in *Drosophila* and mouse models of tauopathy, as well as in human brains affected by Alzheimer’s disease or progressive supranuclear palsy [53-55], a “primary” tauopathy. While evidence in *Drosophila* and HeLa cells suggests that pathogenic forms of tau and over-expression of tau, respectively, cause retrotransposition [54, 56], it is currently unknown if retrotransposons mobilize to a greater degree in the human brain affected by Alzheimer’s disease. We thus analyzed overall singleton insertions and L1 endonuclease (L1en)-mediated singleton insertions in brains lacking tau pathology (Braak 0) and at early (Braak III) and late (Braak V/VI) stages of tau deposition. All three groups featured enrichment of singleton insertions within the centromere when plotted across a meta-chromosome (**Fig. 1E)**. While abundance of general singleton insertions does not differ among Braak stages (**Fig. 1F**), restricting our analysis to L1en-mediated singletons reveals a trending increase in AluY retrotransposition in brains with the highest degree of tau burden (**Fig. 1G, Supplemental Fig. 1C**). The number of Alu and LINE-1 insertions detected per sample is likely an undercount resulting from relatively low sequencing coverage. Our findings from this analysis suggest that key AluY and LINE-1 family members are mobile in cells of the aged human brain and/or their precursor cells and that *de novo* insertion events tend to occur more frequently in the centromere.

### Non-allelic homologous recombination between repetitive elements in the aged human brain

Repetitive element-derived structural variants can arise from NAHR events between homologous high-copy repeats in the genome. NAHR of repetitive elements has been implicated in multiple settings of disease [17, 57, 58], including a recent study utilizing short and long-read DNA sequencing that reports changes in somatic repetitive element-derived NAHR events in neurons isolated from brains affected by Alzheimer’s disease [17].

To delve deeper into structural events linked to repetitive elements, we utilized the TE-rex pipeline to identify singleton repetitive element-derived NAHR events and broadened our analysis to encompass various classes of repetitive elements. CHM13 and GRCh38 genomes were utilized to leverage their respective strengths: assembly completeness (CHM13) vs. more thoroughly documented annotations (GRCh38).

We first identified recombination hotspots across the CHM13 genome by counting NAHR events within 0.1 Mbp windows and normalizing counts by the number of repetitive elements present within the window [17]. To obtain a genome-wide view of NAHR, enrichment values were plotted along a meta-chromosome, which revealed that centromeric/pericentromeric regions exhibit the highest observed normalized enrichment values (**Fig. 2A**), consistent with previous work [17]. We then determined the distribution of NAHR enrichment values for large chromosomal regions, which revealed increased recombination activity primarily in acrocentric chromosome arms, rDNA arrays and centromeric/pericentromeric regions compared to other genomic areas (**Fig. 2B, Supplemental Fig. 2A**). Most recombination events were predicted to result in intrachromosomal rearrangements (**Supplemental Fig. 2B**), consistent with previous findings in neurons.

**Figure 2.**
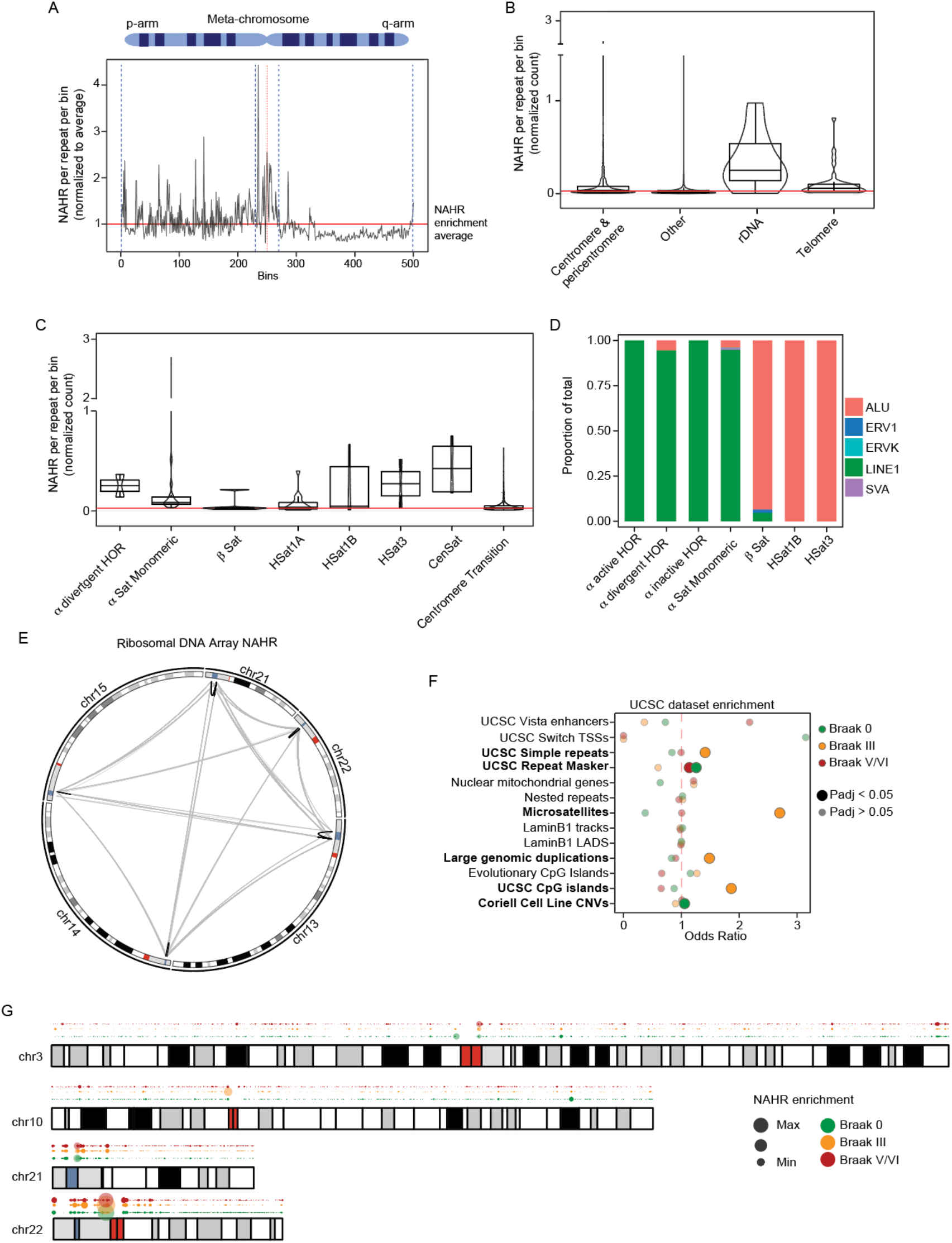
Non-allelic homologous recombination in the aged human brain. A) Meta-chromosome of the rolling mean of the NAHR enrichment values across all autosomes and normalized for chromosome length. B) Boxplots of singleton NAHR enrichment values for different chromosomal structures. C) Boxplots of singleton NAHR enrichment values for different centromeric repeat structures. D) Barplot of transposable element-associated NAHR events among different centromeric repeat structures. E) Circular karyotype map depicting intrachromosomal (gray) and interchromosomal (black) NAHR between rDNA arrays. F) Dotplot of NAHR event enrichment on different UCSC genomic sets based on Braak stage. G) Karyotype map depicting hotspots of NAHR based on Braak stage. Red lines in B, C indicate the average NAHR value across the genome.

Given the high enrichment of NAHR in centromeric and pericentromeric regions, we analyzed NAHR within the functional repeat arrays that compose these regions. NAHR events were detected in HOR repeats (monomeric and divergent), classical human satellite regions, beta satellites, and other centromeric repeat regions (**Fig. 2C**). Interestingly, we find that NAHR HOR regions are composed primarily of L1-L1 recombination events, while NAHR within beta and classical human satellites reflects recombination between Alu elements (**Fig. 2D**). We also detect unique individual regions of enrichment in many peritelomeric and telomeric regions such as the telomeric region of chromosome 17, as well as intergenic segmental duplications such as those present in chromosomes 4 and 6 (**Supplemental File 2**).

Strikingly, 47S rDNA arrays were found to have higher median recombination enrichment compared to other NAHR-prone regions, such as telomeres or the centromere, despite a lower frequency of detected rDNA NAHR events. While the majority of NAHR rDNA events are intrachromosomal, we also identified interchromosomal NAHR events measuring 35 kbp, 45 kbp, and 95 kbp in length (**Fig. 2E, Supplemental Fig. 2B, 2C, 2D**). These events appear to be tandem repeats of the 45 kbp 47S rDNA array, suggesting that duplications and deletions may stem from recombination of full-length rDNA arrays and could underlie variations in rDNA copy number that have been reported in other organisms [59, 60]. While insertion and inversion NAHR rDNA events are marked by Alu-Alu recombination, deletion events appear to stem primarily from LINE-LINE NAHR events, with additional contribution of rRNA-HSA repeats (**Supplemental File 2**).

While the overall frequency of repeat-associated NAHR events does not significantly differ among Braak stages, in agreement with previous analyses of isolated neurons [17] (**Supplemental Fig. 3A-O**), varying values of NAHR enrichment can be discerned in distinct regions of the genome based on Braak stage (examples provided in **Figure 2F, 2G**, full karyotype maps provided in **Supplemental Fig. 4A**). UCSC dataset enrichment analysis further reveals a change in NAHR localization to genomic regions consisting of simple repeats, microsatellites, CpG islands, and annotated segmental duplications at Braak III (**Fig. 2F, 2G**).

**Figure 3.**
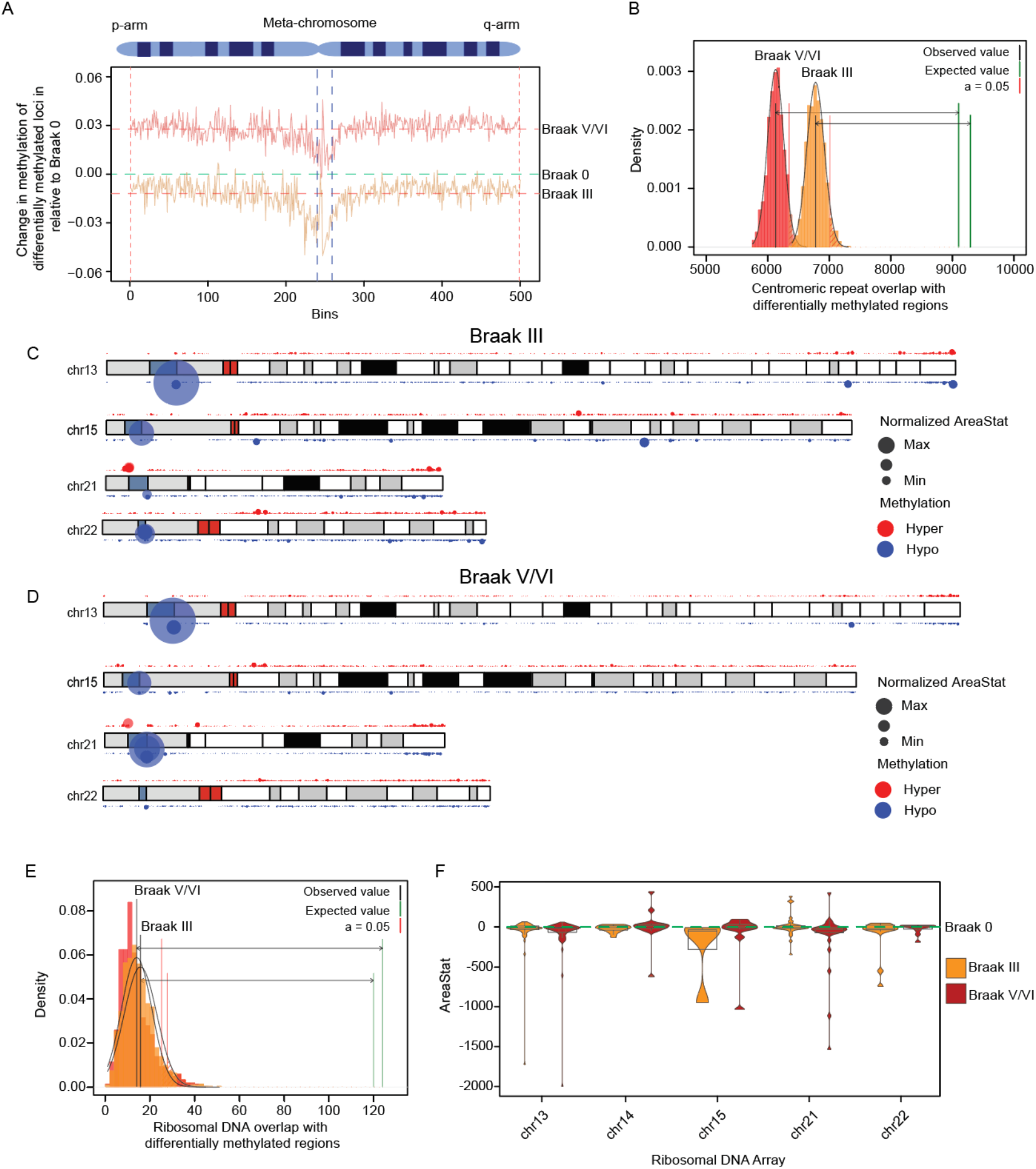
DNA methylation in brains affected by Alzheimer’s disease. A) Meta-chromosome of the rolling mean of the difference in methylation values of differentially methylated regions across all autosomes and normalized for chromosome length for Braak III vs. Braak 0 and Braak V/VI vs. Braak 0. Horizontal dashed lines represent the average change across the whole meta-chromosome for each Braak stage. B) Permutation test of differentially methylated regions localized to the centromere in Braak III vs. Braak 0 and Braak V/VI vs. Braak 0, P=0.0001. C, D) Karyotype of example chromosomes with significant changes in methylation in Braak III and Braak V/VI. E) Permutation test of differentially methylated regions localized to rDNA arrays in Braak III vs. Braak 0 and Braak V/VI vs. Braak 0, P=0.0001. F) Box/violin plot of areaStat values for differentially methylated regions between Braak III vs. Braak 0 and Braak V/VI vs. Braak 0.

**Figure 4.**
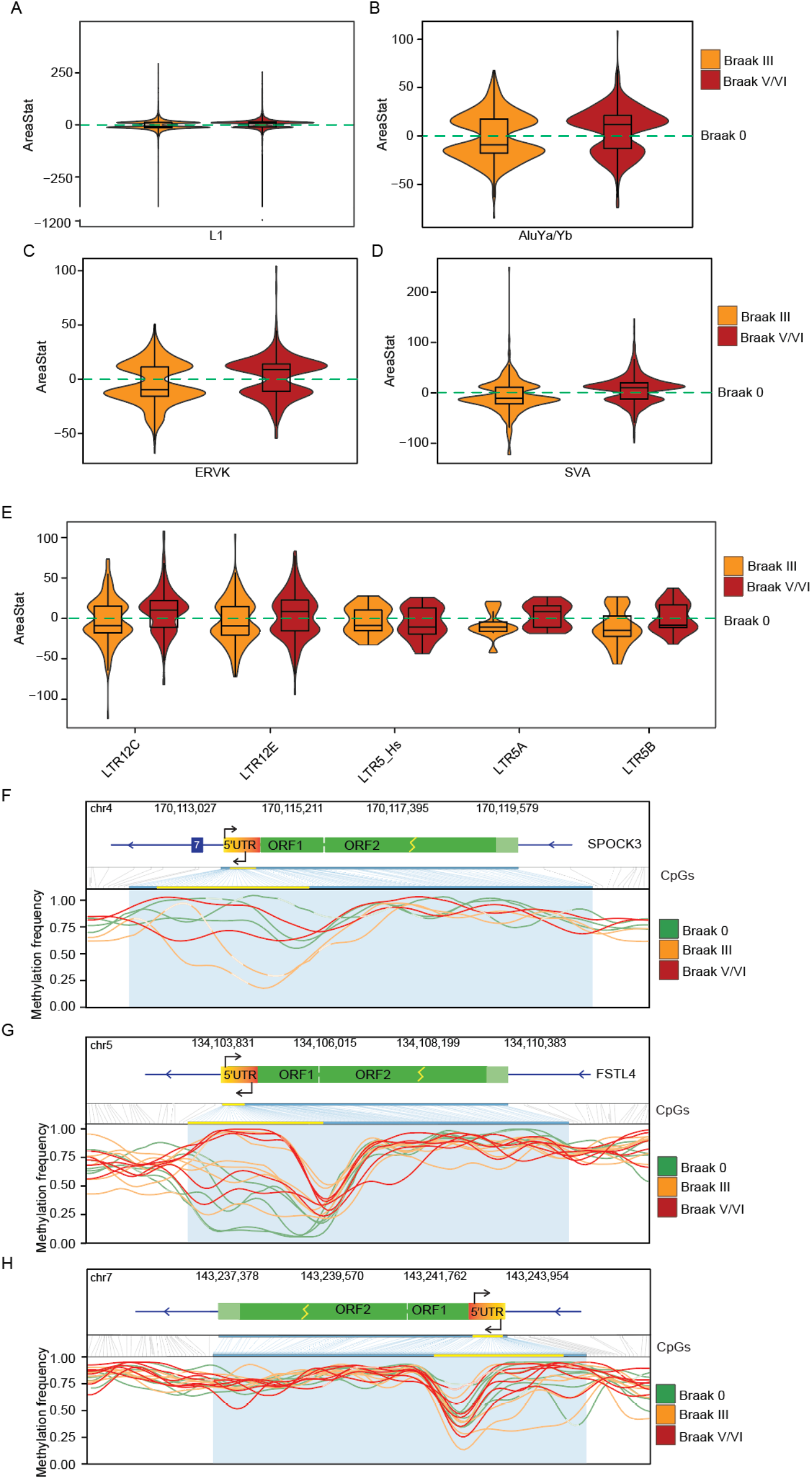
DNA methylation patterns of retrotransposons in Alzheimer’s disease. Box/violin plots of areaStat values for DMRs between Braak III vs. Braak 0 (orange) and Braak V/VI vs. Braak 0 (red) for LINE-1 family (A), AluYa/Yb (B), ERVK subfamily (C), SVA subfamily (D), and LTR sequences (E). F, G, H) Methylation frequency of CpG sites across loci per sample, colored by Braak group. Statically significant changes in CpG methylation detected in promoter regions are highlighted in yellow; yellow lightning bolts denote truncated ORFs. Lighter colors within lines in F and H indicate gaps in sequencing.

### Structural variants in the aged human brain

We next leveraged our dataset to identify complex, repetitive structural variants in the human brain using the more thoroughly annotated GRCh38 genome (**Supplemental Fig. 5A**). Structural variants were most commonly detected in introns and exons (**Supplemental Fig. 5B)**. We find that length distribution of structural variants is highly variable, with insertions having a larger median length compared to other variants (**Supplemental Fig. 5C**). We next utilized SNPeff [61] to predict how these variants may affect gene function. Most structural variants were classified as modifier variants in non-coding genes or intergenic regions or moderate (non-disruptive variants) impact variants (**Supplemental Fig. 5D**). We detect structural variants in genes that are associated with neurodegeneration, including an intronic deletion/inversion in *APP*, a tandem duplication in *ATXN1* [62], and a deletion within *MAPT* (**Supplemental File 3)**.

We next identified non-reference structural variants that fall within the 6,054 dark regions of the human GRCh38 reference genome [18]. Among the 18 brains analyzed, we detect 1,123 insertions in the dark regions of 787 genes, 60 of which occur in dark exons. 1,781 deletions associated with dark regions were detected among the 18 brains analyzed. Interestingly, 23 Alzheimer’s disease risk genes contain dark regions with previously-undocumented structural variants, some of which are in close proximity to SNPs defined as risk variants for neurodegenerative diseases based on the Alzheimer’s Disease Sequencing Project (ADSP) [63] or the European Bioinformatics Institute genome wide association study (EFO ID: EFO_0005772). For example, three dark region-associated insertions and a retrotransposon insertion were detected near or overlapping Alzheimer’s disease-associated SNPs within ATP binding cassette subfamily A member 7 (*ABCA7*) (**Supplemental Fig. 5E**), which encodes an ABC transporter that regulates lipid metabolism [64, 65], amyloid processing and clearance [66-68]. A deletion and an insertion were detected within a dark region of an intronic SVA retrotransposon within vasoactive intestinal peptide receptor 2 (*VIPR2*), near an Alzheimer’s disease-associated SNP that overlaps a deletion present in 10 out of 18 brains analyzed (**Supplemental Fig. 5F**). Similarly, most brains analyzed have a deletion overlapping a dark intronic AluYa5 element within chromodomain helicase DNA binding protein 2 (*CHD2*) in close proximity to an Alzheimer’s disease-associated SNP (**Supplemental Fig. 4G, Supplemental Table 3**). Dark region insertions were also detected near SNPs associated with amyotrophic lateral sclerosis, multiple system atrophy, multiple sclerosis, and spinocerebellar ataxia. Many non-reference insertions and deletions within dark regions were shared among all samples, likely reflecting common circulating variants, while others were only detected in single samples, reflecting less common variants, variants that occurred in the germline, or variants of somatic origin. The overlap of structural variants within dark regions of genes associated with human disease highlights the value of approaches that resolve complex regions of the genome.

We then identified structural variants unique to Braak 0, Braak III or Braak V/VI samples in the CHM13 and GRCh38 reference genomes. To ensure each detected variant is unique to a particular group, we required that the given locus was genotyped in all 18 samples. The number of structural variants detected per sample did not significantly differ between Braak 0, Braak III, and Braak V/VI brains (**Supplemental Fig. 6A, 6B**), nor did structural variant length (**Supplemental Fig. 6C**). These results also hold true when Braak III and V/VI are combined into one group and compared to Braak 0.

### DNA methylation in the aged human brain

We next analyzed DNA methylation patterns (5mC) using the CHM13 and GRCh38 genomes. A sample from an individual with a previous cancer diagnosis was an outlier in terms of 5mC methylation and was removed from subsequent analysis (**Supplemental Fig. 7A**). We focused on CpG sites with a 5mC call in at least three samples, resulting in 24.8 million and 23 million CpG sites for analysis based on the CHM13 and GRCh38 genomes, respectively. This is less than the previously estimated number of CpG sites in the CHM13 (32.28 million) and GRCh38 human genome reference (29.17 million)[69], likely reflecting relatively low coverage sequencing. Our methylation calling approach generally aligns with previous bulk DNA methylation studies of human brain affected by Alzheimer’s disease [32, 70-72].

We initially analyzed changes in differentially methylated regions across the CHM13 genome for Braak III versus Braak 0 and Braak V/VI versus Braak 0 (**Supplemental Fig. 7B, 7C**). We plotted the rolling average of the change in methylation of statistically significant differentially 5mC methylated regions across a meta-chromosome (excluding sex chromosomes) for each comparison. While the average change in methylation was negative in Braak III and positive in Braak V/VI compared to Braak 0 (**Fig. 3A**), some regions share a conserved directional change in Braak III and Braak V/VI stages compared to Braak 0. Centromeric/pericentromeric regions, for example, are demethylated at Braak III and V compared to Braak 0, albeit to varying degrees. Indeed, a permutation test reveals that centromeric methylation is significantly dysregulated in tau-affected brains compared to control (**Fig. 3B, Supplemental Figs. 7D-H**).

To investigate this observation further, we analyzed the area statistic scores of differentially methylated regions. The area statistic (areaStat) reflects the extent of change in a differentially methylated region, considering both its length and the count of CpG sites that undergo methylation alterations. A larger area suggests a higher probability of the region being a genuine differentially methylated region [73]. We determined the distribution of differentially methylated regions across autosomes in Braak III and Braak V/VI cases by plotting areaStat of demethylated regions (DMRs) on a karyotype graph (**Supplemental Fig. 8A, 8B**). Focusing on chromosomes 13, 15, 21, and 22 as examples, we identified shared DMRs between Braak stages based on magnitudes of change in the same or opposite direction (**Fig. 3C, 3D**). Interestingly, these regions overlap with rDNA array model regions of the CHM13 genome. Indeed, a permutation test demonstrated that rDNA regions are enriched for DMRs in Braak III and Braak V/VI compared to Braak 0 (**Fig. 3E**). We detect methylation changes across all five rDNA array regions, with the largest magnitude of disease-associated change being hypomethylation. This is in contrast to a previous observation of hypermethylation in rDNA promoter en-masse in Alzheimer’s disease samples [74]. These findings at Braak III align with previous reports of decreased DNA methylation in cortical tissue of Alzheimer’s disease samples measured by 5mC immunoreactivity [29, 30], although those studies analyzed brains at later Braak stages. Our findings at Braak V/IV are consistent with studies using an array-based approach that reported DNA hypermethylation in Alzheimer’s disease [32-34].

We next analyzed 5mC changes within promoters across Braak stages using the GRCh38 genome. We detect differential methylation of 20,352 unique promoters of 7,852 genes at Braak III compared to Braak 0, while 19,638 promoters of 5,765 genes were found to be differentially methylated at Braak V/VI (**Supplemental File 4**). We observe a subtle change in the bimodal distribution of 5mC modifications in differentially methylated promoters, in line with promoter element MDS (multidimensional scaling) analysis **(Supplemental Fig. 9A, 9B)** [75]. Differentially methylated regions in brains at Braak III have higher densities at 0.25 and 0.75 compared to Braak 0, while brains at Braak V/VI shift towards no methylation (0) or complete methylation (1) (**Supplemental Fig. 9A)**. While promoter methylation differs among Braak 0, Braak III, and Braak V/VI brains, we identify over 7,072 hypomethylated promoters that are unique to brains at Braak III related to pathways regulating phospholipase D, FC gamma receptor mediated phagocytosis, gonadotropin-releasing hormone (GnRH) signaling pathway, Rap1 signaling pathway and actin cytoskeleton regulation (**Supplemental Fig. 9C**). Almost 3,000 hypermethylated promoters are shared between Braak III and Braak V/VI brains, related to pathways involving calcium and oxytocin signaling, focal adhesions, phospholipase D and glutamatergic synapses (**Supplemental Fig. 9C**).

We then focused on DNA methylation within “dark” regions of the genome to fully leverage our long-read approach. We detect 1,150 and 1,333 differentially methylated regions at Braak III versus Braak 0, and Braak V/VI versus Braak 0, respectively. We identified 29 differentially methylated 5’ untranslated regions (UTRs) in Braak stage III and 37 differentially methylated 5’ UTRs in Braak stage V/VI (**Supplemental File 4**). For example, we detect hypomethylation of dark regions in the 5’ UTR of Alzheimer’s disease risk genes *AMY1A* and *FMR1*.

Additionally, we identify four differentially methylated regions in Braak III versus Braak 0 within loci previously identified as Alzheimer’s disease risk genes (*ABACA7, AMY1A, CHRFAM7A, CR1*). Changes in methylation of dark regions in *AMY1A, CHRFAM7A*, and *CR1* were also observed at Braak stage V/VI (**Supplemental File 4**). The detection of a large number of differentially methylated dark regions, including transcription-associated 5’ UTRs, highlights the importance of dark region analyses in epigenetic studies. It is important to note that changes in methylation could represent changes in cell composition between Braak stages rather than (or in addition to) changes in transcriptional control.

### DNA methylation within transposable element sequences of the aged human brain

Consistent with prior observations, we find that methylation levels correlate with the evolutionary age of retrotransposons, with younger elements exhibiting higher methylation frequencies than older elements in the human brain [76-78]. Retrotransposons are heavily methylated in the aged human brain, with a mean methylation value exceeding 0.75, with the exception of HERV-K elements (**Supplemental Fig. 10A, 10B, 10C, 10D, 10E**). Young Alu and LINE retrotransposon family members have generally increased levels of 5mC compared to older Alu and LINE family members, perhaps reflective of the need to repress their transpositional capacities. LTR and HERV families, however, feature less methylation of younger elements.

While previous studies report widespread changes in 5mC at different stages of Alzheimer’s disease in the frontal cortex [32], methylation within dark regions has not been studied in brains of patients with Alzheimer’s disease. An MDS plot of CpG methylation frequency in autosomal repetitive elements revealed no clustering based on sex or Braak stage (**Supplemental Fig. 11A**). While we do not detect differences in 5mC methylation of retrotransposon subfamilies between Braak stages when retrotransposons are grouped *en masse*, similar to previous studies [79, 80] (**Supplemental Fig. 11B-H**), methylation within repetitive regions is clearly dysregulated at the level of individual retrotransposon loci. The distribution of 5mC modification in differentially methylated repetitive elements shifts in opposite directions according to Braak stage, with generally less methylation at Braak III and generally increased methylation at Braak V/VI.

We analyzed the methylation areaStat for select families and subfamilies of retrotransposons to more precisely quantify the degree of differential methylation. We observe large magnitude changes in areaStat for L1 elements, with most of the more significant changes reflective of hypomethylation **(Fig. 4A)**. We separated the L1 family into Human-specific (L1Hs, young) and ancestor-shared (old) L1 subfamilies. While the distribution of L1Hs subfamily methylation is similar among Braak stages, ancestral L1 subfamilies exhibit more hypomethylation at Braak III than V/VI (**Supplemental Fig. 11C**). We observe similar changes in methylation for AluYa/Yb, ERVK, and SVA subfamilies (**Fig. 4B-4D**). Given that Long Terminal Repeat (LTR) elements can function as alternative promoters for genes and as promoters for ERVs, we analyzed methylation patterns of the LTR12 and the young LTR5 families. Specifically, within the LTR12 family, we examined the LTR12C subfamily and its closely-related LTR12E subfamily, as LTR12C is activated in cells treated with epigenetic inhibitors [81]. LTR12C exhibits the most pronounced changes in methylation compared to LTR12E and other LTR subfamilies (**Supplemental File 4**). Similarly, the younger LTR5 family demonstrated significant methylation changes (**Fig. 4E**). We also identified differential methylation in several solo LTR5Hs and Alu elements that are predicted to be regulatory regions based on the GeneHancer database (**Supplemental Table 3**).

We then analyzed changes in methylation of selected differentially methylated non-fragmented (> 5,900 bp) L1Hs retrotransposons on a per-sample basis. Four L1Hs loci (**Supplemental Fig. 12A-E)** feature demethylation at the 5’ UTR, suggesting that these elements may be actively transcribed in the human brain. When comparing among Braak stages, we identify statistically significant hypomethylation in the promoter of individual L1Hs loci such as an intergenic L1-Ta (transcriptionally active) (7q34) and an intronic L1-Ta in *SPOCK3* (4q32.3) at Braak III compared to Braak 0 (**Fig. 4F, G, Supplemental File 4**). Our analysis also reveals examples of LINE hypermethylation at Braak III (**Fig. 4H**). We detect promoter demethylation of an intact L1-Ta element [38] that is reported to be a source of somatic retrotransposition in the developing hippocampus [8, 82]. We identify several differentially methylated LTR5Hs loci in both provirus and solo form at Braak III and V/VI (**Supplemental Fig. 13A, B, Supplemental File 5**), and hypomethylation of 5’ LTR5Hs loci adjacent to proviral HERV-K elements (**Supplementary Fig. 5G, H**), one of which (HERV-K107 (7p22.1)) has been implicated in the pathogenesis of human diseases [83-85]. Overall, our analyses provides novel insights into methylation of individual retroelements and highlights the importance of analyzing such elements at the locus level rather than *en masse*.

## METHODS

### Sample collection

18 postmortem frontal cortex samples were selected to capture equal numbers of Braak stages 0, III and V/VI with an average age of 76.6 years (68 – 92). Samples were Caucasian and were relatively balanced for sex and age with respect to Braak group. Samples were provided by Dr. Dennis Dickson of the Mayo Clinic Brain Bank or acquired from the NIH Neurobiobank.

### DNA processing and read alignment

Approximately 80 mg of tissue was homogenized with a Dounce and then centrifuged through a sucrose gradient to isolate nuclei. High molecular weight DNA was extracted from nuclear isolates using a standard phenol chloroform protocol. Library preparation was performed by the University of California Davis genomics core using the SQK-LSK 110 kit (ONT) with shearing to obtain 10 kbp length reads. Libraries were then multiplexed for three samples per PromethION flow cell and sequenced. Nanopore-derived reads were aligned to the GRCh38 genome (UCSC) and the HS1-CHM13 genome using MiniMap2 [86] (nanopore sequencing specific parameters) to obtain bam files. Mosdepth [87] was used to determine the average coverage per sample.

### Non-reference retrotransposon insertion calling

All bam files were analyzed simultaneously with TLDR (default parameters) to reduce false negative calls in any single sample. This resulted in a retrotransposon file with calls across all 18 samples. Insertions called by TLDR were then filtered to retain insertions with a tandem site duplication and a PASS quality score to eliminate low confidence/quality insertions. We define an L1 endonuclease-like singleton in a post-mitotic cell as a call that is only supported by a single insertion-spanning read (singleton) in a single individual who has at least 10 additional reads lacking the insertion, along with a TSD and L1 endonuclease motif at the insertion site and whose read length was 10Kbp or longer.

### NAHR analyses

Nanopore-derived reads were aligned to both the GRCh38 and CHM13 genomes using the Last alignment software [88]. NAHR events were then called using TE-rex [17] and recommended analysis pipeline to obtain singleton and polymorphic events. Events were only retained if they had a value equal to or less than 1E-4. NAHR enrichment values were calculated by dividing the genome into 100 kbp windows and obtaining the ratio of NAHR events to the total repetitive elements per window.

### Structural variant calling

Individual bam files were subjected to structural variant calling using SNIFFLES2 (default parameters) and subsequently combined into a multi-sample variant call file with the SNIFFLES2 multi-sample calling module. The variant call file was then filtered to retain variants 50 bp or larger. The locations and potential effects of the structural variants in the GRCh38 were determined using snpEFF. Insertion nucleotide sequences were extracted from the variant file and searched against the NCBI nucleotide and RefSeq [89] databases using BLASTn [90] (default parameters) within the Galaxy platform [91]. Hits were retained if the query insertion was covered by 50% or more of a subject sequence in the database. To compare structural variants across Braak groups, we required that the region of a given structural variant was sequenced across all 18 samples. A variant that was only found in samples belonging to a single Braak group was considered unique to that group.

### Normalization and statistical comparison of non-reference retrotransposon insertions and structural variants

TLDR insertions, TE-rex NAHR events, and SNIFFLES2 variants unique to a Braak group were compared by ANOVA. To account for differences in sequencing coverage, insertion counts were normalized by dividing the raw count value by the total number of diploid human genomes sequenced (gbp sequenced divided by 6.4 gbp) (**Supplementary Table 2**).

### Characterization of retrotransposon insertions and structural variants

Coordinates from the TLDR, TE-rex, and SNIFFLES2 output were processed with the R GenomicRanges package [92]. Overlap of insertions and variants with regions (genic/non-genic, dark regions, tandem repeat regions) were determined with AnnotatR [93]. Bedtools [94] was used to identify variants near single nucleotide polymorphisms. Gene graphs were first generated by IGV [95] and then modified for clarity. L1Hs subfamilies were identified by aligning L1Hs loci to the L1.2 (active LINE-1: GenBank accession number M80343) and identifying the L1-Ta (L1Hs-Ta) diagnostic ACA and G nucleotides in the 3’ UTRs. HERV-K open reading frames were determined by aligning HERV-Ks of interest to the consensus HERV-K sequence (DFAM ID: DF0000188). HERV-K non-reference insertions were manually analyzed for coding potential and completeness compared to the full-length HERV-K consensus sequence. To identify other HERV insertions, SNIFFLES2 identified insertions were searched against a custom HERV internal and LTR5 database with sequences from RepBase [96] using BLASTn [90] (default parameters) within the Galaxy platform.

### CpG 5mC methylation analysis

5mC modifications were called using the Nanopolish [97] call-methylation pipeline with single bam files described above on both the GRCh38 and CHM13 genomes. Alignments with a mapping quality score of 20 or more were included in calculating log-likelihood methylation ratios. Nanopolish-generated log-likelihood ratio values were then converted to methylation frequencies with accompanying Nanopolish scripts. Samples were then analyzed via MDS to visualize sample clustering. Sample CTRL-1 had a previous cancer diagnosis and was identified as an outlier; this sample was removed from further analysis. The average methylation frequencies across all samples were annotated as within genes, CpG islands or repeat regions (defined by UCSC RepeatMasker) using annotatR. Methylation graphs of genomic loci were created with methylArtist [38], methylkit [98], and NanomethViz [99].

### Differential methylation analysis

Differentially methylated CpG sites were identified using DSS [73], a tool that models methylation distributions with a beta-binomial model to compute changes in methylation at the loci and regional level, taking sequencing depth into account. Differentially methylated CpG sites were considered significant if they reached a False Discovery Rate (FDR) of 0.05 or less for both the GRCh38 and CHM13 genomes. DSS was then used to identify and aggregate neighboring CpG sites into regions with differential methylation (DMRs) [100]. A sample CpG site methylation frequency was only included if the call was supported by three or more nanopolish called sites. Comparisons were made for Braak III vs. Braak 0 **(Supplemental File 7)** and Braak V/VI vs. Braak 0 **(Supplemental File 8)**. Alu elements (AluYa8b, AluYa5, etc.) and LTR5Hs with the largest changes in methylation between groups were then manually inspected using the GRCh38 GeneHancer Regulatory Elements and the Gene Interaction table [101] within the UCSC genome browser [102] to determine if any overlap occurred. For inclusion in methylation average comparisons of retrotransposons *en masse*, individual loci were required to have 10 or more called CpG sites. To exclude highly fragmented retrotransposon loci, a length threshold depending on the type of retrotransposon was implemented: HERV-K (> 6,000 bp), LTR5_Hs/A/B (> 900 bp), LINE1-Hs (> 5,900 bp), AluYa5/Yb8 (> 280 bp) and SVA-E/F (> 1,000 bp) [38]. Other repetitive elements such as centromeric satellite regions were required to have 10 or more CpG sites called within a given region. For analysis of DMRs in promoter regions, a promoter region was only counted once in each direction, even if there are multiple differentially methylated regions within the region.

### Enrichment analysis and statistical tests

A permutation test (regioneR function permTest [103]) was used to determine if there are more differentially methylated regions within centromeres and rDNA arrays than what would be expected by chance. The mean change in methylation in differentially methylated regions across a meta-chromosome was calculated following previously described methods [104, 105]. Gene lists were analyzed for KEGG pathway [106] enrichment using ClusterProfiler [107]. All graphs were generated with R packages ggplot2 [108], TreeMap or adapted from IGV [95]. Lola [109] was used for genomic region enrichment with UCSC datasets. KaryotypeR was used for karyotype graphs [110].

## DISCUSSION

Retrotransposon insertions, repetitive element-associated NAHR, complex structural variants, and DNA modifications within dark/repetitive regions have long been understudied due to the technical difficulty of their detection with traditional sequencing methods. The emergence of long-read sequencing technologies such as Oxford Nanopore, associated bioinformatic tools, and more complete human genome references allow us to overcome previous limitations. Here, we detect genomic and epigenomic changes in the aged human brain, with an unexpected predominance of events within centromeric/pericentromeric regions and rDNA arrays.

The centromere is highly repetitive region that has thus been historically overlooked in most genomic studies [111]. We detect striking genomic changes within the centromere across brain samples. We discovered a significant enrichment of retrotransposon insertion singletons, particularly Alu elements, within centromeric alpha satellite HOR arrays. Analyses of NAHR events at the centromere revealed a distinct pattern related to transposable element loci: L1-L1 NAHR events were found to predominantly occur within alpha satellite monomeric repeats, while Alu-Alu recombination events are enriched in pericentromeric and acrocentromeric regions. In the context of Alzheimer’s disease, we detect a clear depletion in methylation at the centromere in both early and late stages of disease. This finding adds a new dimension to previous studies reporting decondensation of constitutive heterochromatin, which is enriched at the centromere, in laboratory models of tauopathy and in neurons isolated from postmortem human Alzheimer’s disease brain [112-114].

Our analyses point toward rDNA arrays as a second hotspot for genomic variation in the human brain, in agreement with previous analyses in other tissues [115, 116]. We detect numerous Alu-Alu NAHR events occurring within and between rDNA arrays across different chromosomes, in line with a report of enriched break points near Alu sequences in rDNA [117]. Interestingly, we find that rDNA is demethylated at early and late stages of tau pathology compared to control. Future studies should investigate the potential interplay between rDNA NAHR, methylation loss, and the potential formation of circular rDNA structures, as observed in other organisms and human cell lines [59, 118]. Observed genomic changes in rDNA arrays may underlie reported changes in translation rate in settings of tauopathy [119-122] or the overall fidelity of the translatome.

The elevation of retrotransposon transcripts in Alzheimer’s disease and progressive supranuclear palsy, along with increased transposition in a *Drosophila* model of tauopathy suggest that transposable elements are derepressed and may mobilize in human Alzheimer’s disease [53-55, 123-126]. While our analysis did not reveal a significant difference in singleton/L1en-mediated insertions between brains at Braak 0, III, and V/VI, we detect a clear trend of increasing AluYa5 insertion with elevated tau pathology. Our nanopore analyses further revealed clear differences in DNA methylation at specific Alu and LTR5Hs loci (previously implicated as alternative promoters [127-130]) based on Braak stage. Several L1Hs loci feature altered promoter methylation specific to disease stage (albeit with high intergroup variation), indicating that they may serve as source loci for active L1 elements in Alzheimer’s disease [131]. Our analyses clearly highlight the value of epigenetic analyses of individual retrotransposon loci compared to grouping such elements by family.

While previous analyses of structural variants in brain cells, often relying on DNA copy number and DNA content, have revealed alterations that accumulate with age and in settings of disease, these approaches can miss complex variants entirely, hindering a comprehensive understanding [132]. This has resulted in a lack of information regarding structural variants and/or pathogenic retrotransposon insertions that are in linkage disequilibrium with SNPs. Our analyses reveal several insertions in a dark region composing a variable nucleotide repeat expansion that is highly correlated with a common high-penetrant risk SNP (rs3764650) for Alzheimer’s disease in the lipid transporter gene *ABCA7* [133, 134]. We also detect overlap between an Alzheimer’s disease risk SNP (rs115550680) and a polymorphic AluYa8 element, and identified several challenging-to-detect variants within other known risk genes. Our analyses likely reflect the tip of the iceberg regarding potential contributions of genetic variants within dark regions to disease; future studies leveraging deeper long read sequencing and/or targeted sequencing methods such as Cas9, PCR, and hybridization capture in a larger set of individuals will enable comprehensive population-level variant analysis [135].

Taken together, our analyses reveal enrichment of repetitive element-derived genomic alterations within centromeric, pericentromeric, and rDNA regions in cells of the aged human brain, suggesting that these areas of the human genome are particularly susceptible to instability. As long-read sequencing technologies continue to advance and gain broader adoption, they will undoubtably increasingly uncover clinically-significant genomic and epigenetic variations associated with human disorders.

## DATA AVAILABILITY

All sequencing data will be made publicly available in the NCBI SRA database (PRJNA1083482) upon acceptance of the manuscript.

## CODE AVAILABILITY

Example code used to generate data will be deposited in the Zenodo repository.

## ACKNOWLEDGEMENTS

The Texas Advanced Computing Center (TACC) at The University of Texas at Austin provided HPC resources that contributed to the analysis and research results reported within this paper. URL: http://www.tacc.utexas.edu. DNA sequencing and library preparation were carried out at the DNA Technologies and Expression Analysis Cores at the UC Davis Genome Center, supported by NIH Shared Instrumentation Grant 1S10OD010786-01. Human post-mortem frontal cortex tissues were acquired from the NIH Neurobiobank or Mayo Clinic Brain Bank.

## ETHICS DECLARATIONS

No conflicts of interest are declared.

