## Supplemental Information for "Nanopore Long-Read Sequencing Unveils Genomic Disruptions in Alzheimer’s Disease"

### SUPPLEMENTAL FIGURES

**Supplemental Figure 1 | Non-reference genome retrotransposon insertions in the aged human brain, CHM13 genome**

**Supplemental Figure 2 | Non-allelic homologous recombination in the aged human brain, CHM13 genome**

**Supplemental Figure 3 | NAHR based on Braak stage, CHM13 and GRCh38 genomes**

**Supplemental Figure 4 | Non-allelic homologous recombination based on Braak stage, CHM13 genome**

**Supplemental Figure 5 | Non-reference structural variants in the aged human brain, GRCh38 genome**

**Supplemental Figure 6 | Non-reference structural variants in the aged human brain, CHM13 and GRCh38 genomes**

**Supplemental Figure 7 | Differential methylation analysis based on Braak stage, CHM13 genome**

**Supplemental Figure 8 | Differential methylation analysis based on Braak stage, CHM13 genome**

**Supplemental Figure 9 | Differential promoter methylation analysis based on Braak stage, GRCh38 genome**

**Supplemental Figure 10 | Differential retrotransposon methylation analysis, CHM13 genome**

**Supplemental Figure 11 | Differential retrotransposon methylation analysis based on Braak stage, CHM13 genome**

**Supplemental Figure 12 | Methylation profiles of select full-length L1Hs retrotransposons, CHM13 genome**

**Supplemental Figure 13 | Methylation analysis of select HERVK retrotransposons, CHM13 genome**

### SUPPLEMENTAL TABLES

**Supplemental Table 1 | Demographics of brain samples**

**Supplemental Table 2 | Nanopore sequencing metrics**

**Supplemental Table 3 | Retrotransposon loci methylation metrics and predicted gene associations, GRCh38 genome**

### SUPPLEMENTAL FILES

**Supplemental File 1 | Non-reference genome retrotransposon insertion calls**
Sheet 1 - Main output file of TLDR annotated with supplemental information
Sheet 2 - Genic locations of TLDR insertions

**Supplemental File 2 | TE-rex NAHR calls**
Sheet 1 - Main output file of TE-rex combined VCF file converted into table format, CHM13 genome

Sheet 2 - Main output file of TE-rex combined VCF file converted into table format, GRCh38 genome

**Supplemental File 3 | Non-reference structural variant tables, GRCh38 genome**

Sheet 1 - Main output file of SNIFFLES2 combined VCF file converted into table format
Sheet 2 - Blast results to identify insertions
Sheet 3 - Pseudogenized gene insertions, insertions and deletions identified in dark regions

**Supplemental File 4 | Differential methylation of promoter regions and dark regions, GRCh38 genome**

Sheet 1 - Differential methylation of promoter regions, Braak III vs Braak 0
Sheet 2 - Differential methylation of promoter regions, Braak V/VI vs Braak 0
Sheet 3 - Differential methylation of dark regions, Braak III vs Braak 0
Sheet 4 - Differential methylation of dark regions, Braak V/VI vs Braak 0

**Supplemental File 5 | Differentially methylated repetitive regions, CHM13 genome**

Sheet 1 - Differential methylation of repetitive regions, Braak III vs Braak 0 and Braak V/VI vs Braak 0

**Supplemental File 6 | Raw output of TLDR analysis, CHM13 genome**

**Supplemental File 7 | Insertion calls for non-reference genome structural variants**

Sheet 1 - Raw VCF output of SNIFFLES2 analysis, CHM13 genome
Sheet 2 - Raw VCF output of SNIFFLES2 analysis, GRCh38 genome

**Supplemental File 8 | Raw output of differentially methylated loci at Braak III vs. Braak 0, CHM13 genome**

**Supplemental File 9 | Raw output of differentially methylated loci at Braak V/VI vs. Braak 0, CHM13 genome**

### SUPPLEMENTAL FIGURES

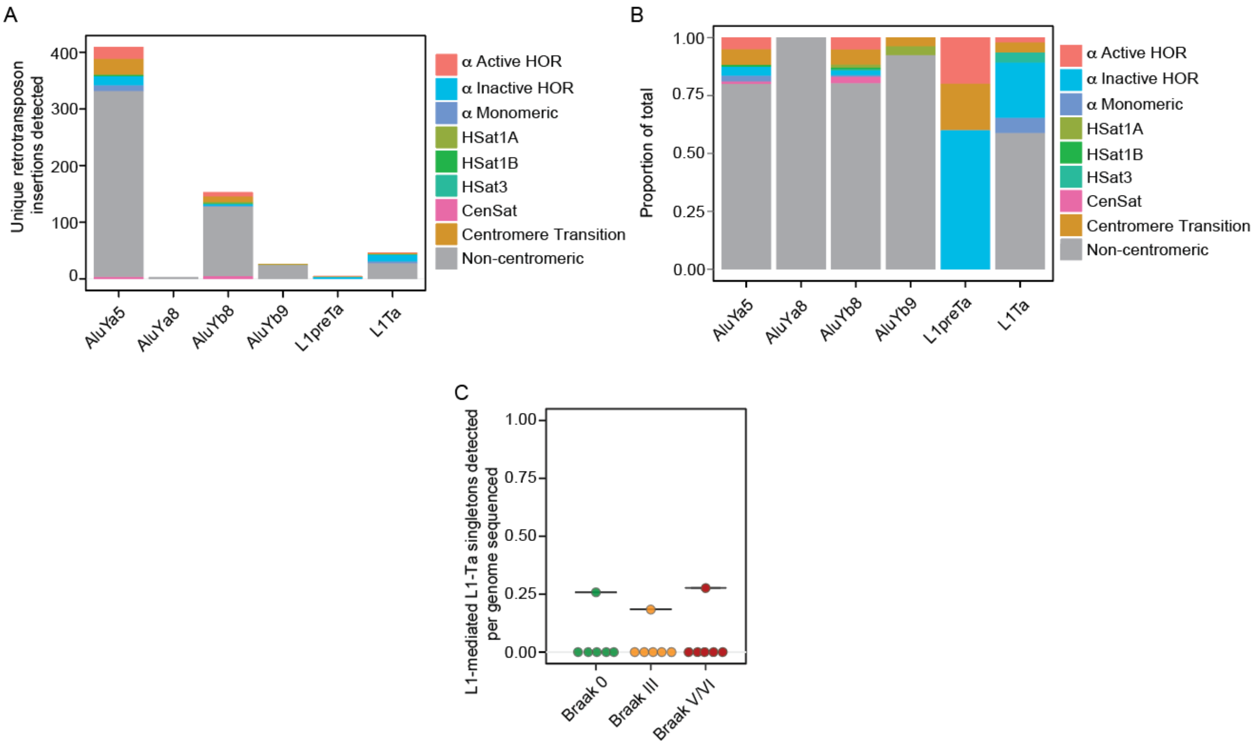


**Supplemental Figure 1 | Non-reference genome retrotransposon insertions in the aged human brain, CHM13 genome**
A) Bar plot of retrotransposon insertion events unique to a single sample with respect to regions of the centromere. B) Bar plot of the proportion of centromeric regions in which sample-specific retrotransposon insertions occur. C) Box plots of L1en-mediated singleton L1-Ta insertion counts normalized to sequencing depth.

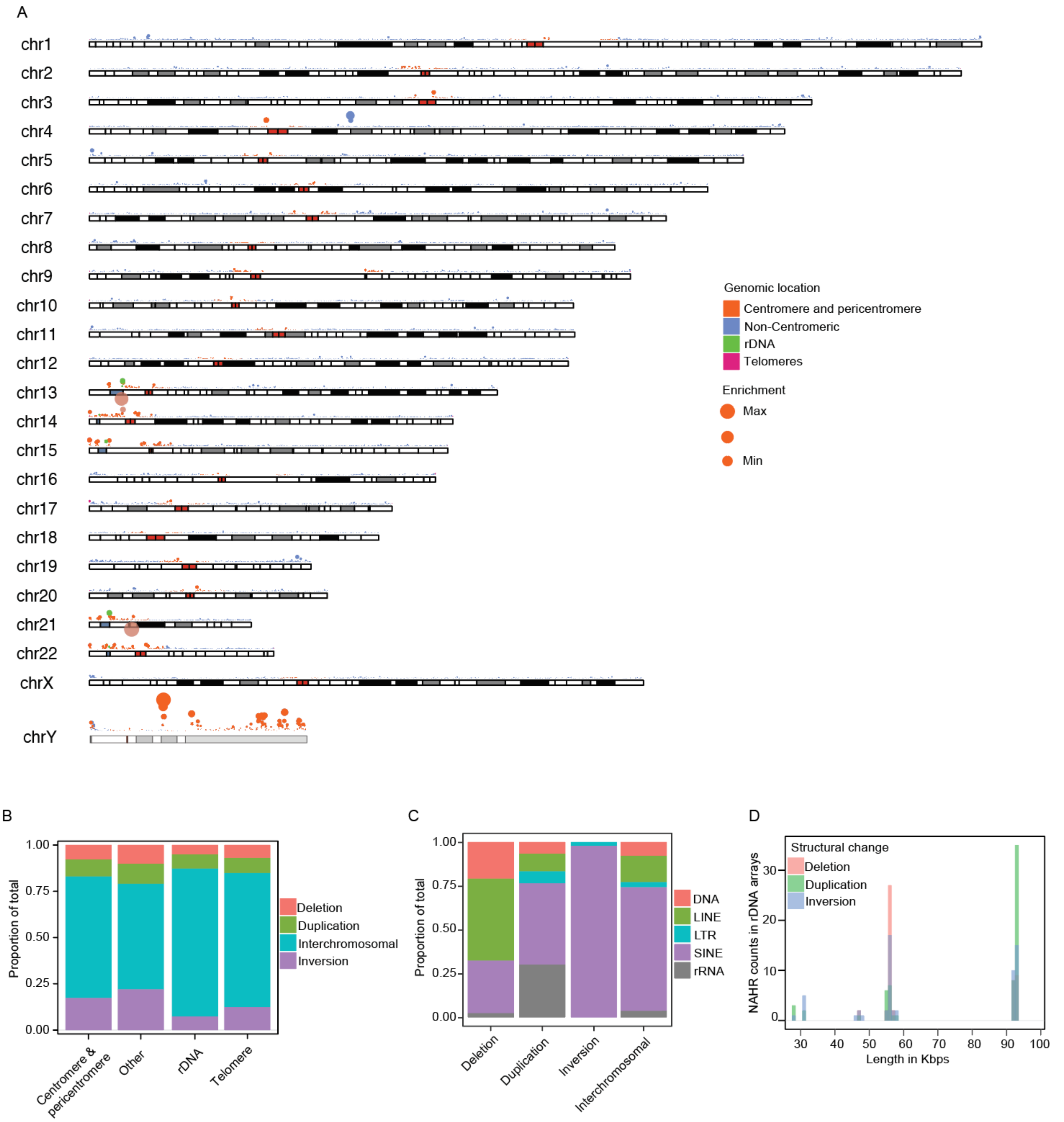


**Supplemental Figure 2 | Non-allelic homologous recombination in the aged human brain, CHM13 genome**

A) Karyoplot of NAHR enrichment across all 18 samples based on the CHM13 genome. B) Bar plot of the proportion of NAHR event-derived structural variants per genomic region. C) Bar plot of the proportion of retrotransposon family NAHR events per structural variant. D) Histogram of intrachromosomal NAHR structural variants in rDNA arrays.

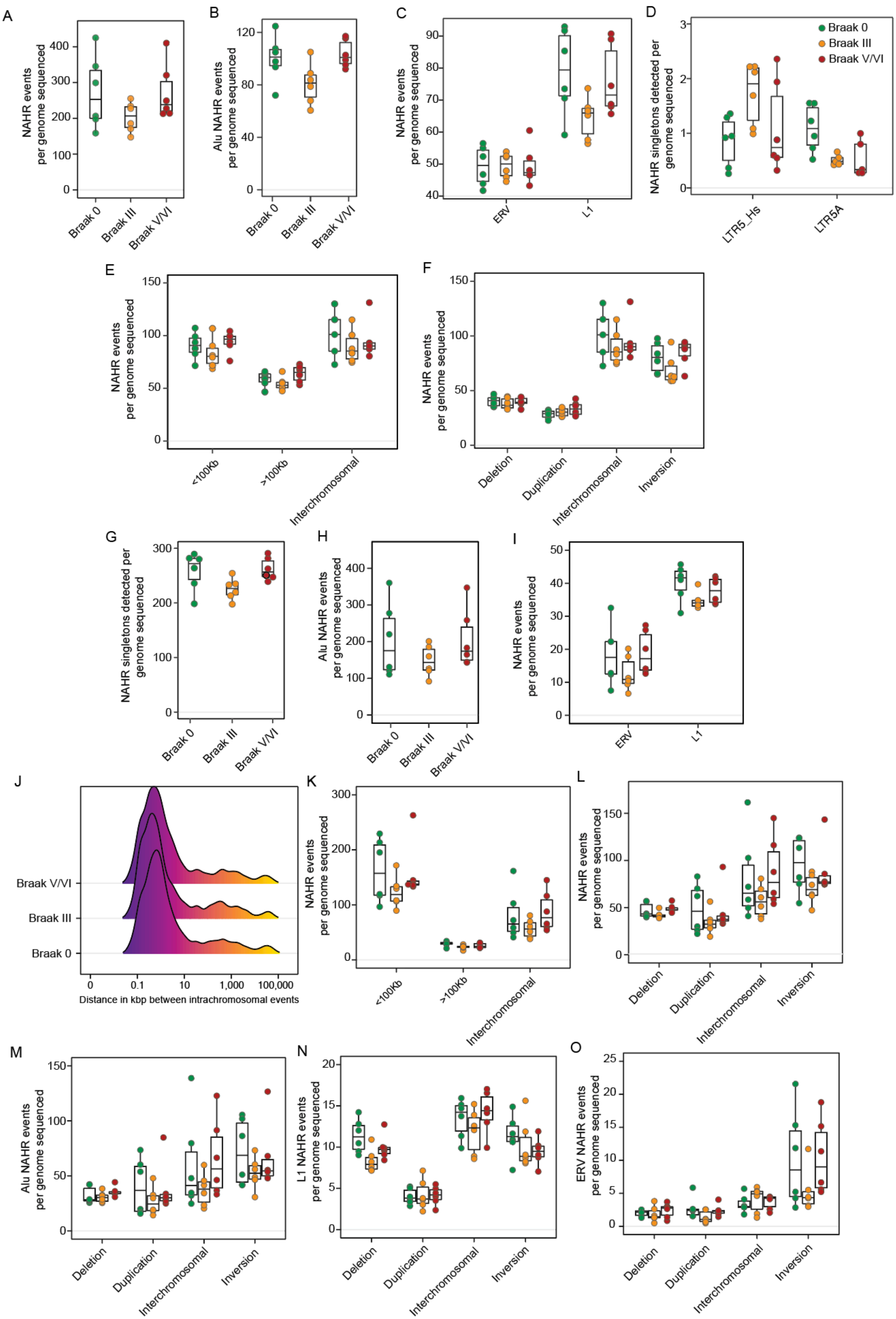

A
NAHR events per genome sequenced
Braak 0
Braak III
Braak V/VI
B
Alu NAHR events per genome sequenced
C
NAHR events per genome sequenced
ERV
L1
D
NAHR singletons detected per genome sequenced
LTR5_Hs
LTR5A
E
NAHR events per genome sequenced
<100Kb
>100Kb
Interchromosomal
F
NAHR events per genome sequenced
Deletion
Duplication
Interchromosomal
Inversion
G
NAHR singletons detected per genome sequenced
H
Alu NAHR events per genome sequenced
I
NAHR events per genome sequenced
J
Distance in kbp between intrachromosomal events
K
NAHR events per genome sequenced
L
NAHR events per genome sequenced
M
Alu NAHR events per genome sequenced
N
L1 NAHR events per genome sequenced
O
ERV NAHR events per genome sequenced

**Supplemental Figure 3 | Analysis of NAHR based on Braak stage, CHM13 and GRCh38 genomes**

A) Box plots of all normalized NAHR singleton counts. B) Box plots of all Alu NAHR singleton counts normalized to sequencing depth. C) Box plots of all normalized ERV, L1 NAHR singletons. D) Density plot of intrachromosomal structural variant length by Braak stage. D) Box plots of normalized LTR5_Hs and LTR5A NAHR singleton counts. E) Box plots of all normalized NAHR singleton counts by length of intrachromosomal events (lesser or greater than 100 Kbp). F) Box plots of all normalized NAHR singleton grouped by structural variant. G) Box plots of all normalized NAHR singleton counts. H) Box plots of all normalized Alu NAHR singletons. I) Box plots of all normalized ERV, L1 NAHR singleton counts. J) Density plot of intrachromosomal structural variant length by Braak stage. K) Box plots of all normalized NAHR singleton counts by length of intrachromosomal events (lesser or greater than 100 Kbp). Box plots of normalized L) NAHR, M) Alu, N) ERV and O) L1 NAHR singleton counts grouped by structural variant. CHM13 genome was used in A-I, GRCh38 genome was used in L-O.

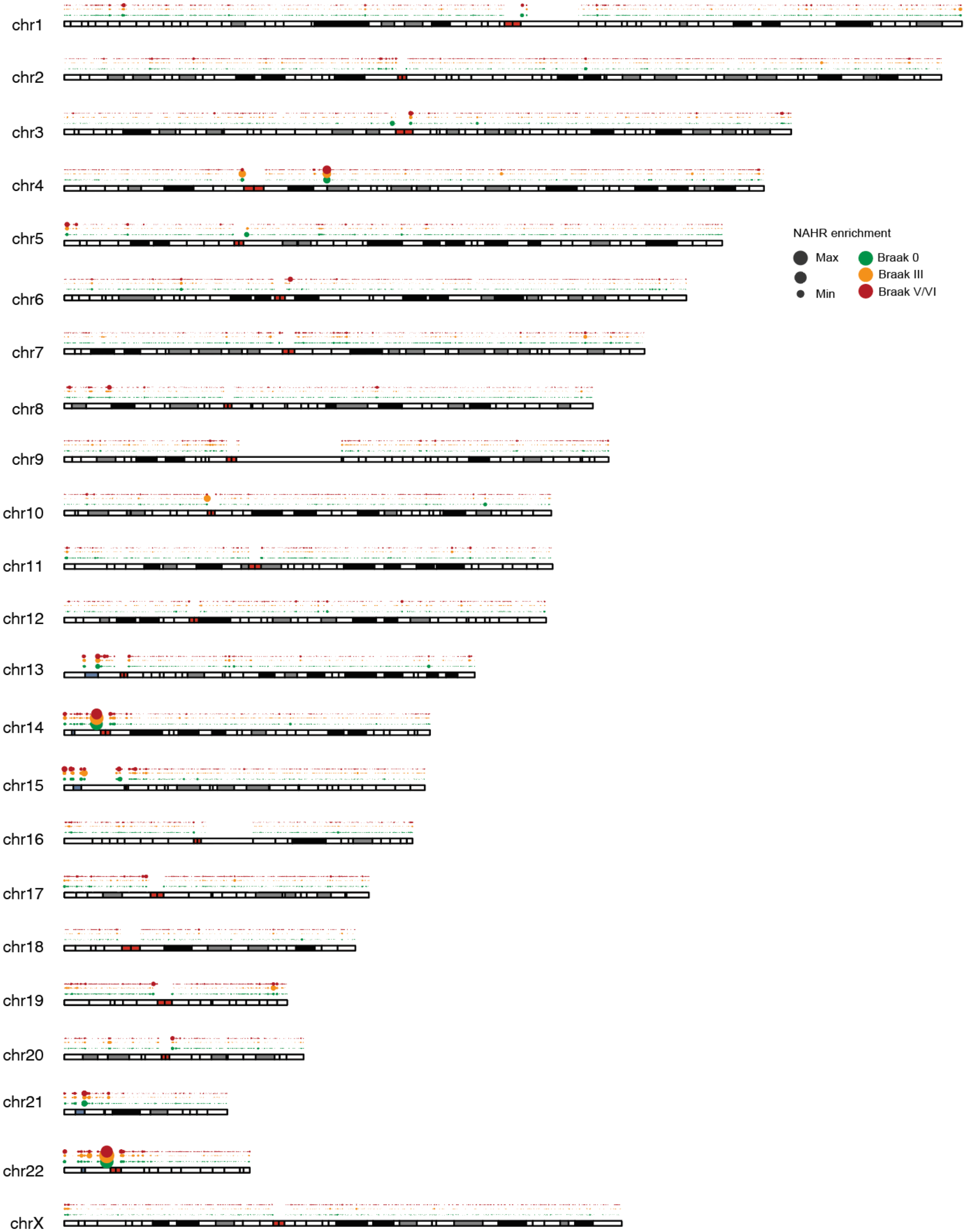


**Supplemental Figure 4 | Non-allelic homologous recombination based on Braak stage, CHM13 genome**
Karyotype plot of NAHR enrichment values pooled by Braak stage.

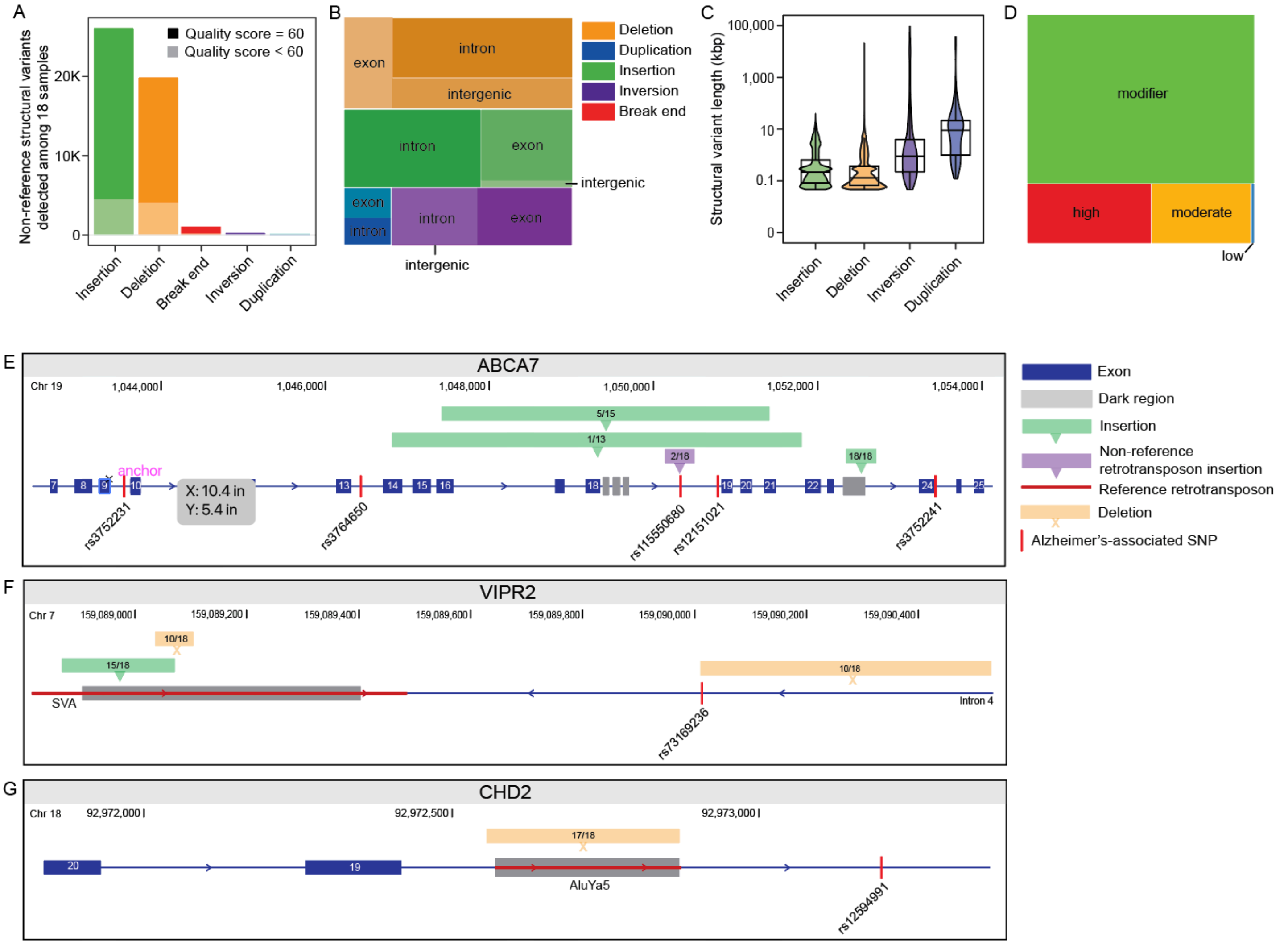


**Supplemental Figure 5 | Non-reference structural variants in the aged human brain, GRCh38 genome**

A) Total counts of all non-reference structural variants detected. Dark shading represents variants with the highest quality score (60), while lighter shading represents lower scores. B) Treemap depicting respective location of structural variant according to gene structure. C) Violin plot of structural variant lengths between 500 bp and 100 mbp. D) Treemap of predicted effects of structural variants on gene function. E) Total structural variants per brain sample detected by Braak stage, normalized by sequencing depth. F) Violin plot of structural variant length (within the range 500 bp – 1 mbp) for structural variant length. E, F, G) Novel insertions and deletions detected in close proximity to Alzheimer's disease associated SNPs. Ratios indicate the proportion of human brains carrying the given variant.

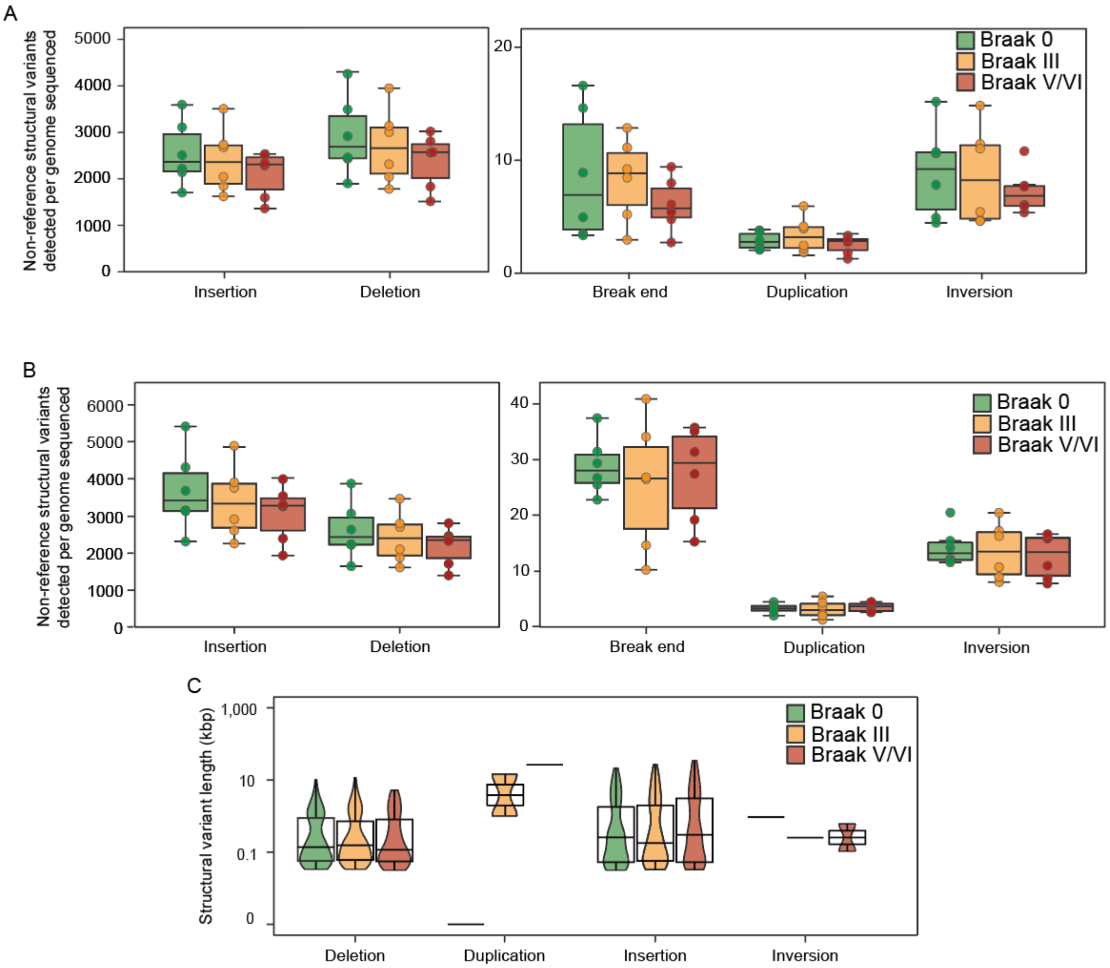


**Supplemental Figure 6 | Non-reference structural variants in the aged human brain, CHM13 and GRCh38 genomes**

A) Boxplots of all non-reference structural variants insertion counts for different structural variants normalized to sequencing depth in the CHM13 genome. B) Total structural variants per brain sample detected by Braak stage, normalized by sequencing depth using the GRCh38 genome. C) Violin plot of structural variant length (within the range 500 bp – 1 mbp) for structural variant length using the GRCh38 genome.

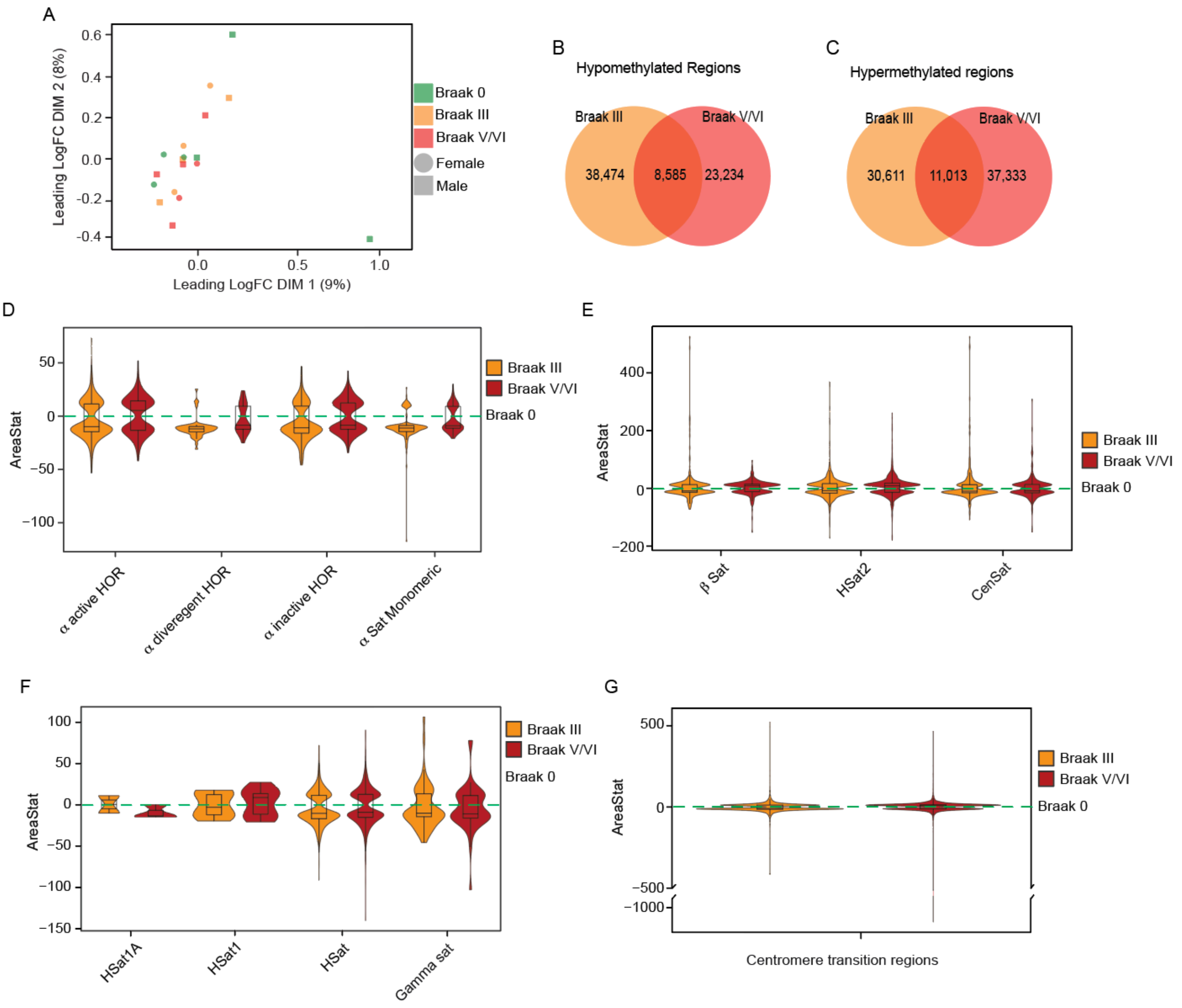


**Supplemental Figure 7 | Differential methylation analysis based on Braak stage, CHM13 genome**

A) MDS plot of likelihood methylation ratio (LMR) of autosomal CpG sites. B, C) Venn diagram of differentially methylated regions shared or unique to Braak III or Braak V/VI. D, E, F, G) Box/violin plots of areaStat values for DMRs between Braak stage III vs. Braak 0 and Braak stage V/VI vs. Braak 0 for centromeric functional repeat regions.

A

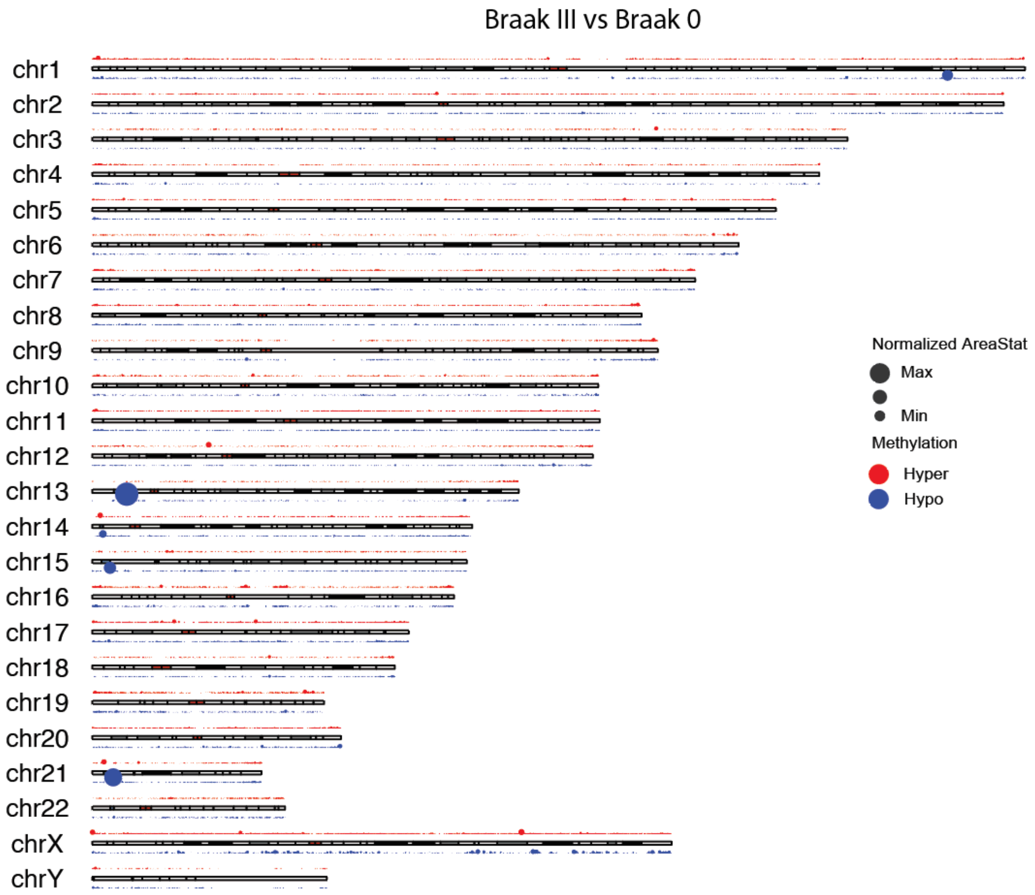
Braak III vs Braak 0
chr1
chr2
chr3
chr4
chr5
chr6
chr7
chr8
chr9
chr10
chr11
chr12
chr13
chr14
chr15
chr16
chr17
chr18
chr19
chr20
chr21
chr22
chrX
chrY
Normalized AreaStat
Max
Min
Methylation
Hyper
Hypo

B

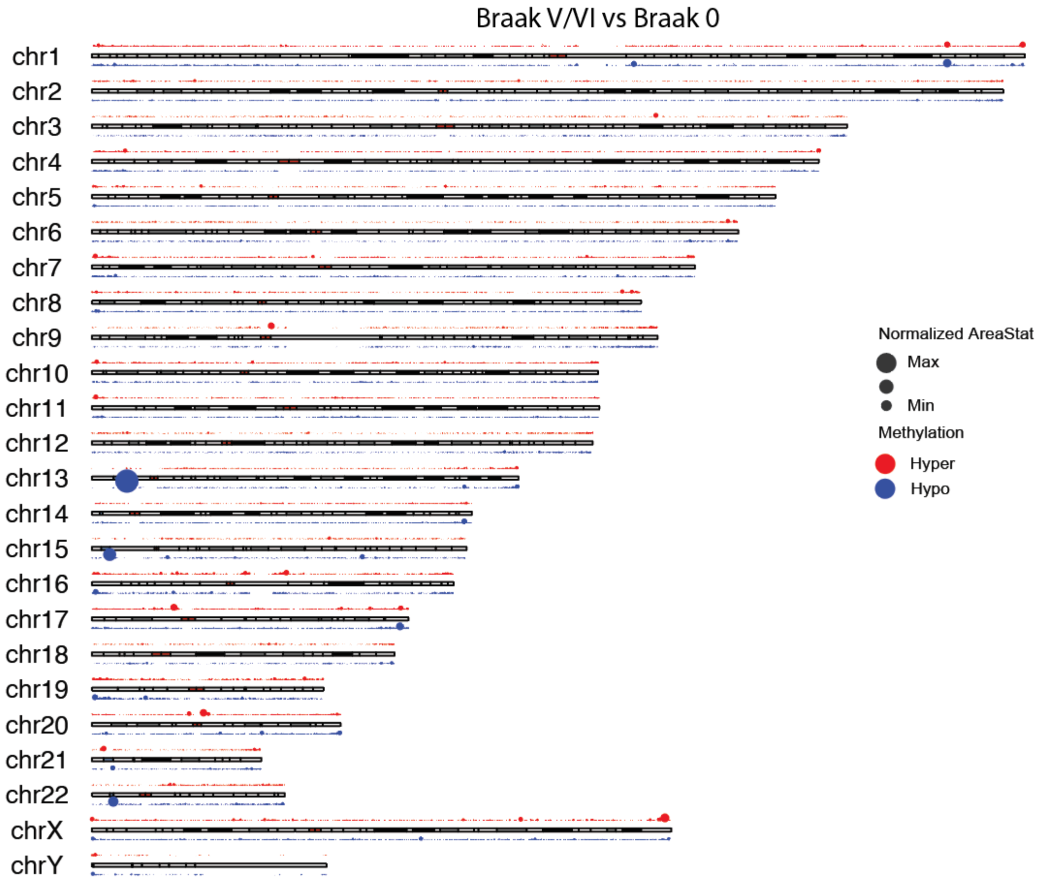
Braak V/VI vs Braak 0
chr1
chr2
chr3
chr4
chr5
chr6
chr7
chr8
chr9
chr10
chr11
chr12
chr13
chr14
chr15
chr16
chr17
chr18
chr19
chr20
chr21
chr22
chrX
chrY
Normalized AreaStat
Max
Min
Methylation
Hyper
Hypo

**Supplemental Figure 8 | Differential methylation analysis based on Braak stage, CHM13 genome**

Karyotype plot of differentially methylated region enrichment in Braak III vs. Braak 0 (A) and Braak V/VI vs. Braak 0 (B).

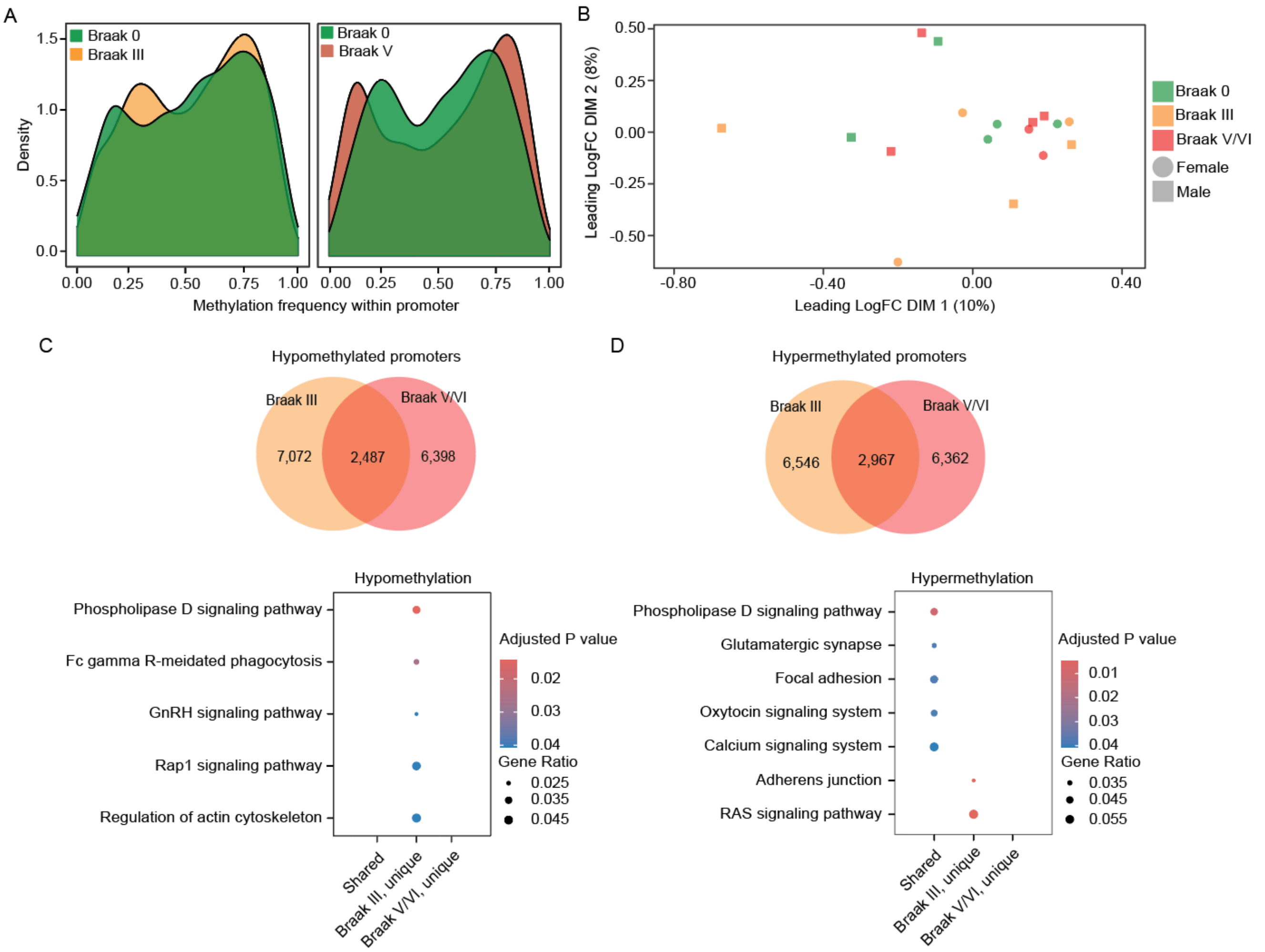


**Supplemental Figure 9 | Differential methylation analysis of promoters based on Braak stage, GRCh38 genome**

A) Density of differentially methylated regions within promoters (Y-axis) at mean methylation frequency in Braak III vs. Braak 0 and Braak V/VI vs. Braak 0 (X-axis). B) MDS plot of likelihood methylation ratios of promotor region CpG sites. C, D) Venn diagram of promoter methylation and respective comparison of functional KEGG profiles of promoters that are shared or unique to Braak III or Braak V/VI.

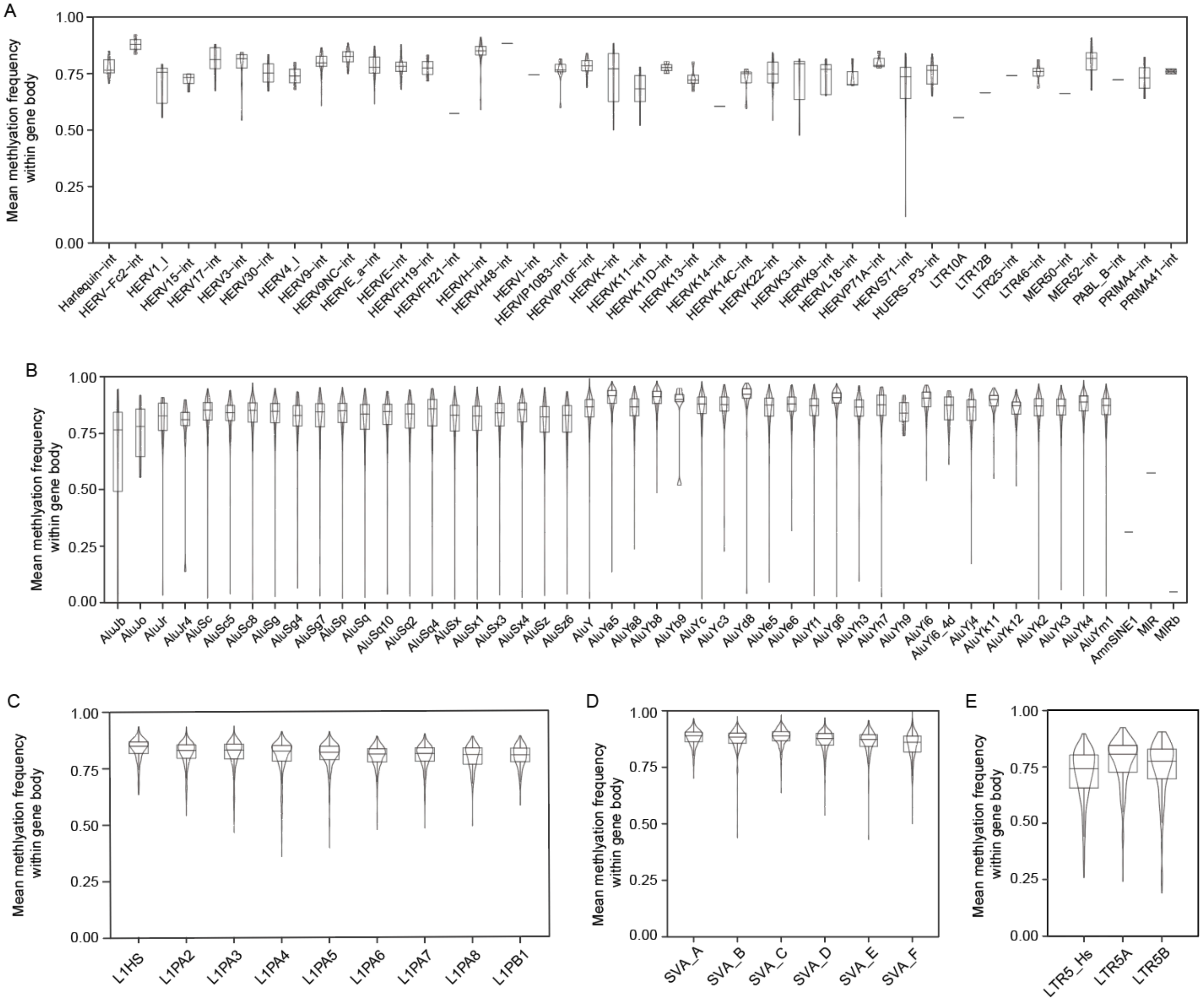


**Supplemental Figure 10 | Methylation analysis of retrotransposon family member, CHM13 genome**

Violin and box plots of the mean methylation frequency within retrotransposon with at least 10 CpG site calls for all samples combined and a minimum length of 5 kbp in ERV elements (A), 290 bp in Alu elements (B), 5.9 kbp in L1 elements (C), 1000 bp in SVA elements (D), and 900 bp in LTR5 elements (E).

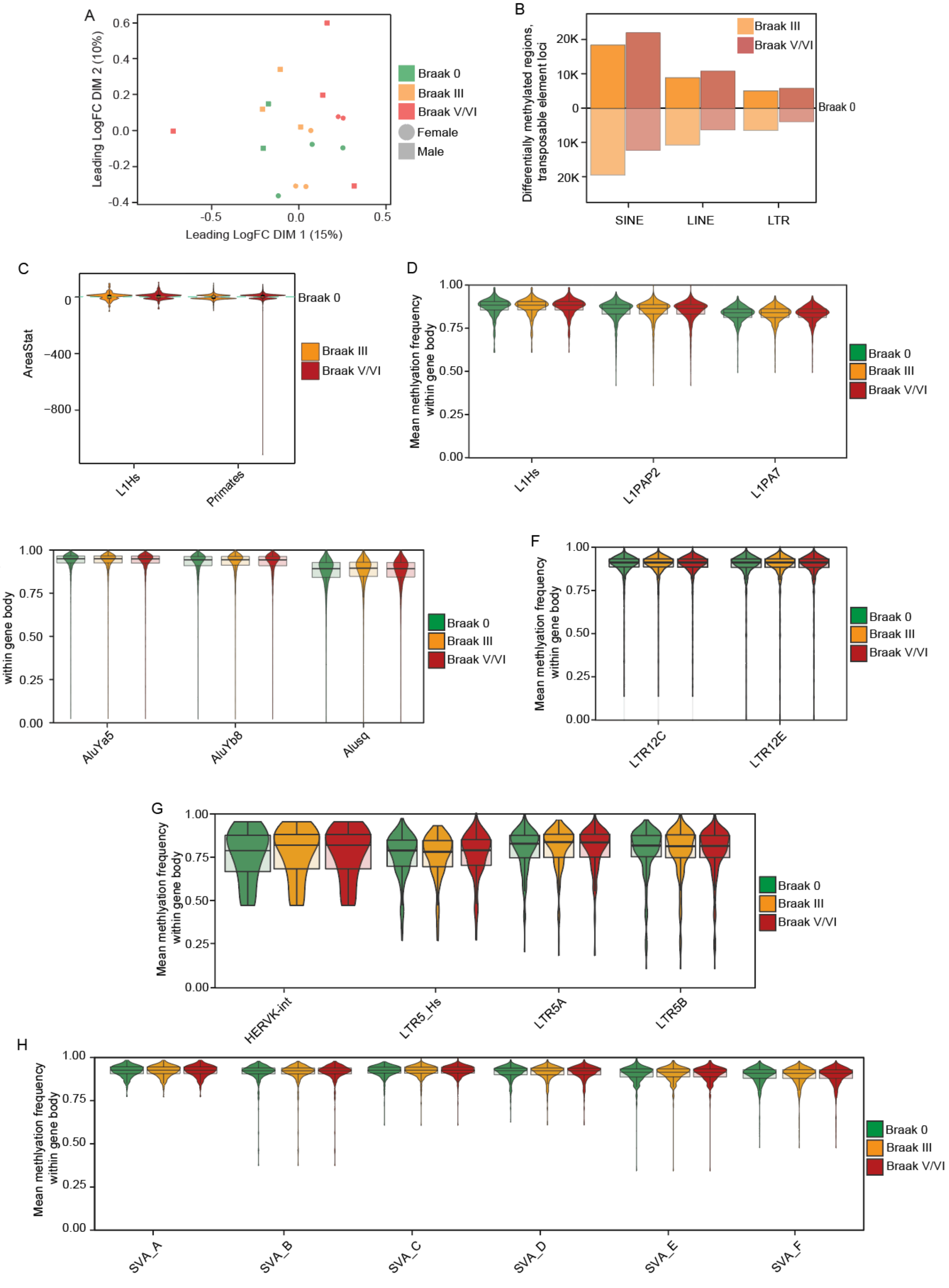


**Supplemental Figure 11 | Methylation analysis of transposable elements based on Braak stage, CHM13 genome**
A) MDS plot of LMR of autosomal CpG sites within repetitive elements. B) Bar plot of hypermethylated and hypomethylated repetitive regions in Braak III vs. Braak 0 and Braak V/VI vs. Braak 0. C) Box/violin plots of areaStat values for DMRs between Braak stage III vs. Braak 0 (orange) and Braak stage V/VI vs. Braak 0 (red) for L1Hs and older L1s. Violin and box plots of the mean methylation frequency

within gene body of L1 (D), Alu (E), LTRs (F), HERV-K/LTR (G) and SVA (H) sequences based on Braak stage for loci with at least 10 CpG site calls with a minimum length of 5.9 kbp in L1, 900 bp in LTR5Hs, 1000 bp in SVA family and 290 bp in AluYa5/Yb8.

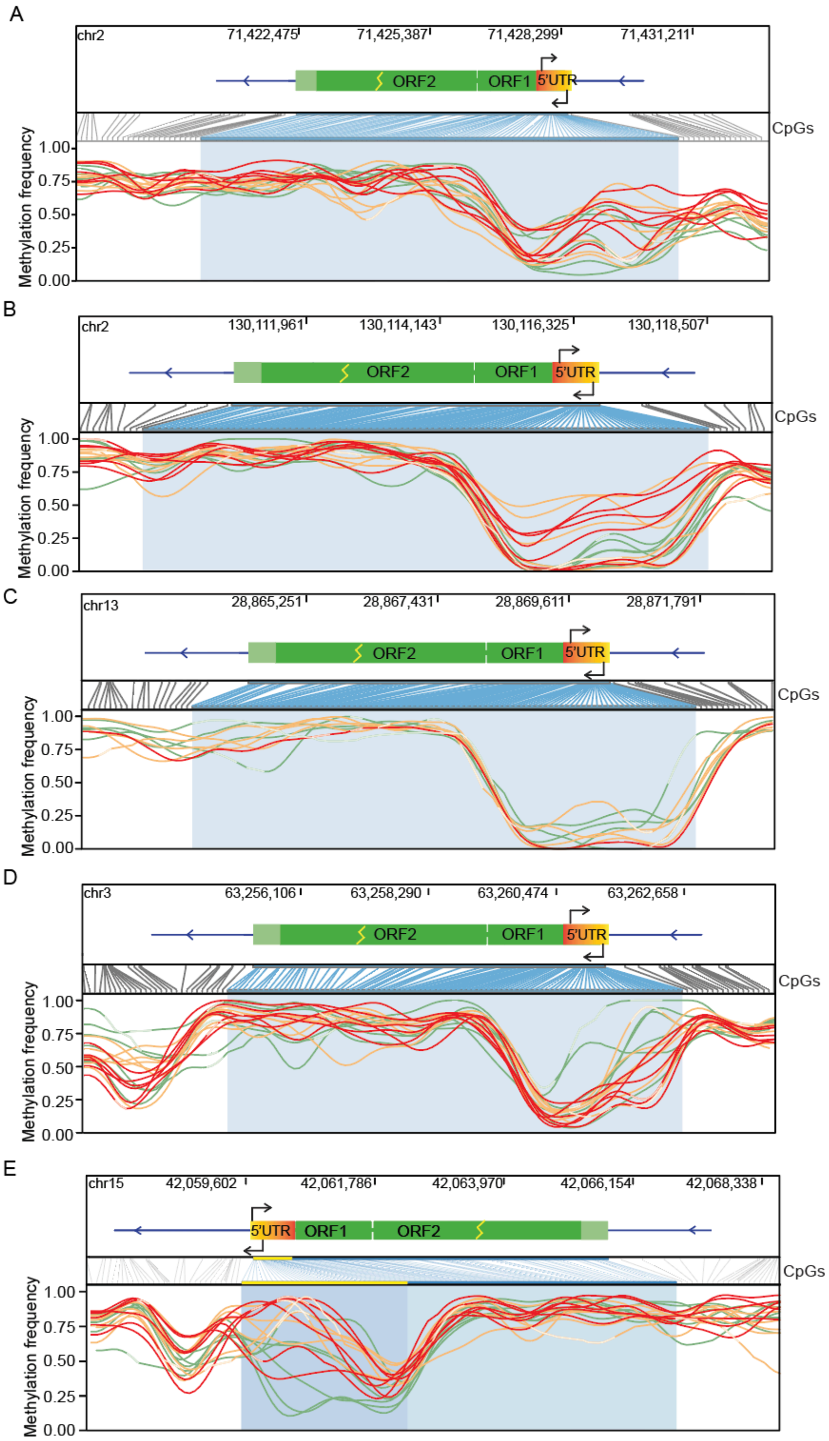


**Supplemental Figure 12 | Methylation profiles of select full-length L1Hs retrotransposons, CHM13 genome**
A, B, C, D, E) Methylation profile of L1Hs retrotransposon loci by sample colored by Braak stage. The yellow bar represents regions identified as differentially methylated based on DSS.

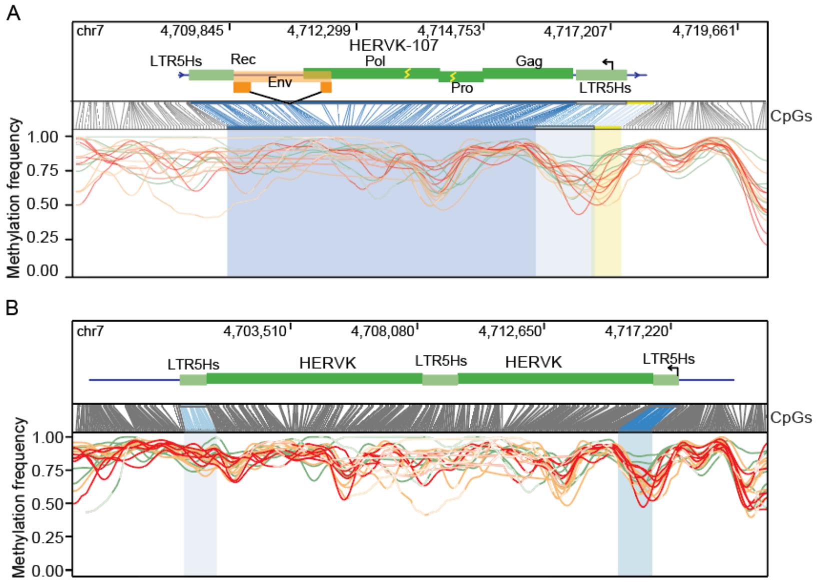


**Supplemental Figure 13 | Methylation analysis of select HERVK retrotransposons, CHM13 genome**

A, B) Methylation profile of HERVK retrotransposon loci by sample colored by Braak stage. The yellow bar represents regions identified as differentially methylated by DSS.

### SUPPLEMENTARY TABLES

| Sample type | Sample name | Braak stage | Thal stage | Age | Sex | PMI | Race |
|---|---|---|---|---|---|---|---|
| Braak 0 | CTRL-1 | 0 | Unknown | 84 | Male | Unknown | Unknown |
| Braak 0 | CTRL-2 | 0 | Unknown | 83 | Male | 18 | Caucasian |
| Braak 0 | CTRL-3 | 0 | Unknown | 68 | Female | Unknown | Unknown |
| Braak 0 | CTRL-4 | 0 | Unknown | 79 | Male | 10 | Caucasian |
| Braak 0 | CTRL-5 | 0 | Unknown | 68 | Female | Unknown | Unknown |
| Braak 0 | CTRL-6 | 0 | Unknown | 74 | Female | 3 | Caucasian |
| Braak III | Braak_III_1 | 2.5 | 0 | 88 | Male | Unknown | Caucasian |
| Braak III | Braak_III_2 | 3 | 0 | 92 | Female | Unknown | Caucasian |
| Braak III | Braak_III_3 | 3 | 0 | 79 | Female | 8 | Caucasian |
| Braak III | Braak_III_4 | 3 | 2 | 75 | Male | 12 | Caucasian |
| Braak III | Braak_III_5 | 3.5 | 0 | 85 | Female | 24 | Caucasian |
| Braak III | Braak_III_6 | 3 | 2 | 75 | Male | Unknown | Caucasian |
| Braak V/VI | Braak_V-VI_1 | 6 | Unknown | 71 | Male | Unknown | Unknown |
| Braak V/VI | Braak_V-VI_2 | 6 | Unknown | 67 | Female | 12 | Caucasian |
| Braak V/VI | Braak_V-VI_3 | 5 | Unknown | 79 | Male | 13 | Caucasian |
| Braak V/VI | Braak_V-VI_4 | 5 | Unknown | 70 | Male | Unknown | Unknown |
| Braak V/VI | Braak_V-VI_5 | 6 | Unknown | 73 | Female | 5 | Caucasian |
| Braak V/VI | Braak_V-VI_6 | 6 | Unknown | 68 | Male | 5 | Caucasian |

**Supplemental Table 1 | Demographics of brain samples**
PMI = Postmortem interval in hours.

| **Sample type** | **Sample name** | **Mean read quality** | **Total reads** | **Read length n50** | **Total gbp** | **Genome coverage** |
|---|---|---|---|---|---|---|
| Braak 0 | CTRL-1 | 11.6 | 1,023,179 | 30,505 | 14.19 | 4.09 |
| Braak 0 | CTRL-2 | 11.9 | 1,627,426 | 30,006 | 18.24 | 5.30 |
| Braak 0 | CTRL-3 | 11.5 | 6,478,290 | 11,670 | 35.20 | 10.00 |
| Braak 0 | CTRL-4 | 11.8 | 2,047,730 | 28,548 | 21.61 | 6.32 |
| Braak 0 | CTRL-5 | 11.8 | 3,840,393 | 21,898 | 25.73 | 7.43 |
| Braak 0 | CTRL-6 | 11.8 | 3,638,404 | 25,346 | 25.78 | 7.51 |
| Braak III | Braak_III_1 | 11.8 | 1,455,635 | 27,275 | 15.69 | 4.57 |
| Braak III | Braak_III_2 | 11.8 | 4,185,578 | 16,613 | 27.67 | 8.02 |
| Braak III | Braak_III_3 | 11.7 | 2,443,785 | 28,230 | 35.66 | 10.50 |
| Braak III | Braak_III_4 | 11.5 | 2,733,026 | 23,740 | 21.07 | 6.11 |
| Braak III | Braak_III_5 | 11.7 | 1,937,159 | 25,730 | 20.02 | 5.88 |
| Braak III | Braak_III_6 | 11.6 | 2,014,256 | 25,769 | 30.55 | 8.93 |
| Braak V/VI | Braak_V-VI_1 | 11.7 | 1,504,113 | 28,758 | 19.63 | 5.75 |
| Braak V/VI | Braak_V-VI_2 | 11.9 | 8,720,004 | 9,668 | 34.08 | 9.74 |
| Braak V/VI | Braak_V-VI_3 | 11.7 | 6,542,225 | 17,291 | 41.64 | 12.12 |
| Braak V/VI | Braak_V-VI_4 | 11.8 | 3,226,405 | 16,556 | 24.18 | 7.01 |
| Braak V/VI | Braak_V-VI_5 | 11.9 | 3,583,034 | 19,083 | 24.58 | 7.05 |
| Braak V/VI | Braak_V-VI_6 | 11.8 | 1,903,934 | 31,580 | 22.55 | 6.59 |

**Supplemental Table 2 | Nanopore sequencing metrics**

Table of Promethion nanopore sequencing metrics per sample. Total reads = total number of reads sequenced. Read length n50 = read length n50 in kbp. Total gbp = total amount of DNA sequenced per sample. Genome coverage = sequencing coverage of sample.

| Name | Coordinates | Stage | CpGs | Control | Braak | Δ 5mc | GeneHancer genes |
|---|---|---|---|---|---|---|---|
| AluYa5 | chr11:102601093-102601457 | Braak III | 13 | 0.81 | 0.6 | -0.21 | MMP7, MMP20 |
| AluYa5 | chr2:20598293-20598970 | Braak III | 17 | 0.53 | 0.7 | 0.17 | Hs13BP, RHOB, GDF7 |
| LTR5_Hs | chr11:63530136-63530437 | Braak III | 15 | 0.52 | 0.32 | -0.2 | GH11J063530, RARRESE3 |
| LTR5_Hs | chr8:144795817-144796432 | Braak III | 10 | 0.35 | 0.18 | -0.17 | ZNF34, RPL8, ZNF517 |
| LTR5_Hs | chr11:63529265-63530678 | Braak V/VI | 15 | 0.62 | 0.45 | -0.18 | GH11J063530, RARRESE3 |

**Supplemental Table 3 | Retrotransposon loci methylation metrics and predicted gene associations, GRCh38 genome**
Table of retrotransposon loci with the largest changes of methylation in Braak III or Braak V/VI. Coordinates = coordinates in GRCh38 reference genome. Stage = Braak stage at which methylation change occurs. CpGs = number of CpG sites present in the differentially methylated region. Control = mean methylation frequency at differentially methylated region for Braak 0. Braak = mean methylation frequency at differentially methylated region for respective Braak group. Δ 5mc =change in methylation. GeneHancer genes = list of genes of which associated retrotransposon are predicted to interact with through an enhancer-based function via the GeneHancer track in the UCSC genome browser.
